# Evolutionary repair reveals an unexpected role of the tRNA modification m^1^G37 in aminoacylation

**DOI:** 10.1101/2021.07.14.452415

**Authors:** Ben E. Clifton, Muhammad Aiman Fariz, Gen-Ichiro Uechi, Paola Laurino

## Abstract

The tRNA modification m^1^G37, which is introduced by the tRNA methyltransferase TrmD, is thought to be essential for growth in bacteria due to its role in suppressing translational frameshift errors at proline codons. However, because bacteria can tolerate high levels of mistranslation, it is unclear why loss of m^1^G37 is not tolerated. Here, we addressed this question by performing experimental evolution of *trmD* mutant strains of *E. coli*. Surprisingly, *trmD* mutant strains were viable even if the m^1^G37 modification was completely abolished, and showed rapid recovery of growth rate, mainly *via* tandem duplication or coding mutations in the proline-tRNA ligase gene *proS*. Growth assays and *in vitro* aminoacylation assays showed that G37-unmodified tRNA^Pro^ is aminoacylated less efficiently than m^1^G37-modified tRNA^Pro^, and that growth of *trmD* mutant strains can be largely restored by single mutations in *proS* that restore aminoacylation of G37-unmodified tRNA^Pro^. These results show that inefficient aminoacylation of tRNA^Pro^ is the main reason for growth defects observed in *trmD* mutant strains and that the ProRS enzyme may act as a gatekeeper of translational accuracy, preventing the use of error-prone unmodified tRNA^Pro^ in protein translation. Our work shows the utility of experimental evolution for uncovering the hidden functions of essential genes and has implications for the development of antibiotics targeting TrmD.

## Introduction

Post-transcriptional chemical modification of tRNA controls the stability, folding, and decoding properties of tRNA molecules and is essential for accurate and efficient protein translation (1, 2). *N*^1^-methylation of guanine 37 (m^1^G37), which is immediately adjacent to the anticodon on the 3’ side, is a tRNA modification that is particularly important for cell growth and is believed to be essential in bacteria (Fig. 1a) (3–5). In bacteria, the SpoU-TrmD (SPOUT) family tRNA methyltransferase TrmD is responsible for introducing the m^1^G37 modification into tRNAs containing the ^36^GG^37^ sequence motif, which includes three proline tRNAs (all isoacceptors), one arginine tRNA (CCG isoacceptor), and three leucine tRNAs (CAG, GAG and UAG isoacceptors) in the case of *E. coli* (4, 6, 7). It has long been known that temperature-sensitive mutations in *trmD* that reduce the level of m^1^G37 modification cause severe growth defects and have a frameshift suppressor effect, increasing the rate of +1 translational frameshifting at Pro codons, which led to the hypothesis that the role of TrmD and the m^1^G37 modification in bacteria is to ensure that the correct reading frame is maintained during protein translation (8). Indeed, frameshift errors in protein translation are highly deleterious, resulting in synthesis of truncated, misfolded, and inactive protein, which could explain why TrmD is essential for cell survival. Based on these observations, it is widely accepted that the primary function of TrmD is to prevent frameshift errors in translation (2, 4, 5, 9, 10). Recent work has provided a detailed view of the mechanism for frameshift errors induced by m^1^G37-deficient tRNA and the system-wide effects of m^1^G37 deficiency at the cellular level (11–14).

**Figure 1.**
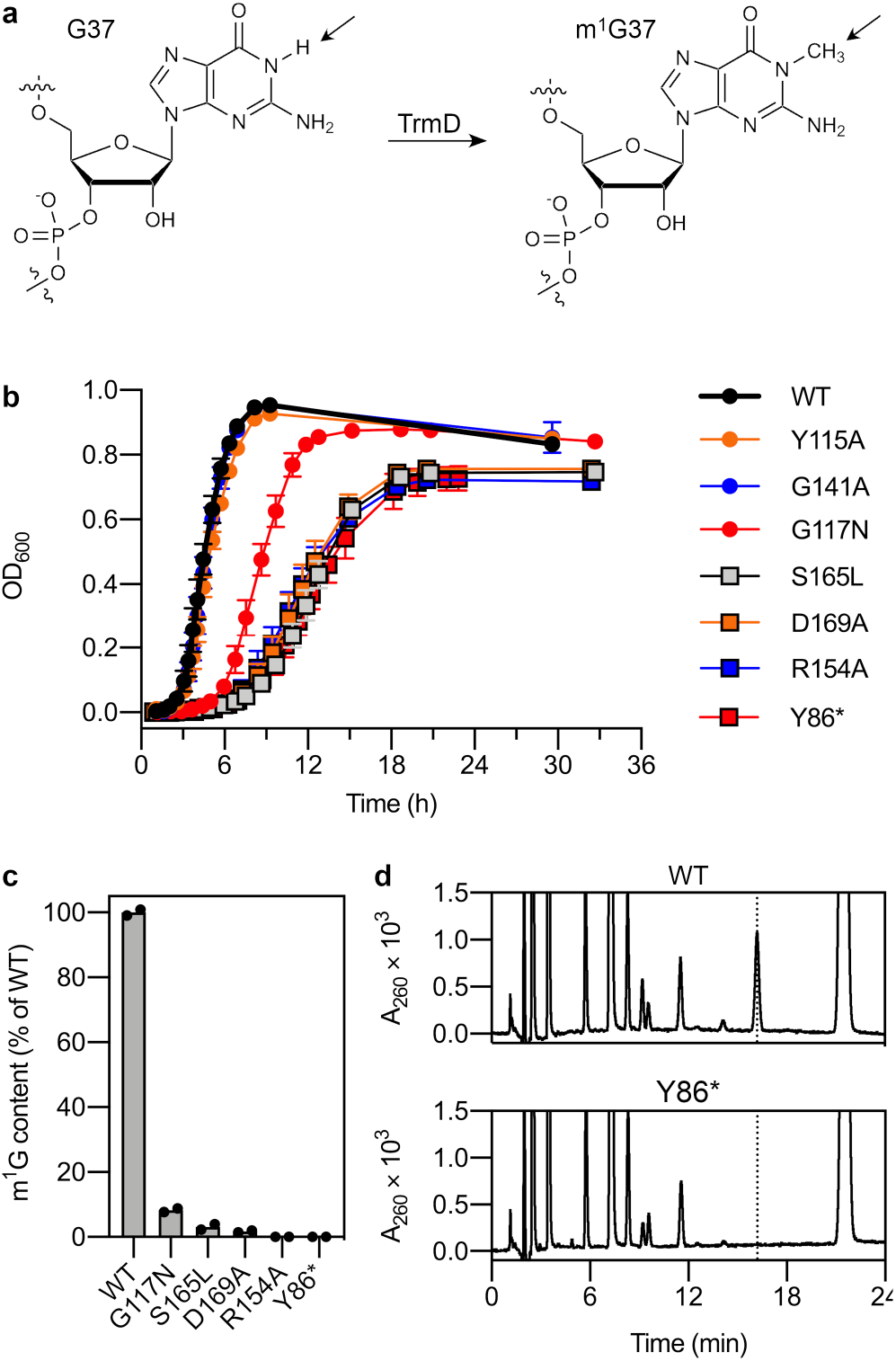
Characterization of*E. coli* strains with mutations in *trmD*. **a**, *N*^1^-methylation of guanosine 37 catalyzed by TrmD. The arrows indicate the position of the modification. **b**, Growth assays of *trmD* mutant strains in LB medium at 37 °C. Data represent mean ± s.d.: WT, Y115A, *n* = 9 colonies; G141A, *n* = 5 colonies; G117N, R154A, Y86*, *n* = 4 colonies (three technical replicates per colony); S165L, D169A, *n* = 3 colonies (three technical replicates per colony). **c,** m^1^G content in tRNA^Pro^ (CGG) purified from *trmD* mutant strains, determined by UPLC. m^1^G content was estimated by comparing the peak area ratio of m^1^G to pseudouridine in the A_260_ chromatogram between the WT and mutant strains. *n* = 2 biological replicates. **d,** Separation of nucleosides in digested tRNA^Pro^ (CGG) by UPLC. UV chromatograms (260 nm detection) are shown for tRNA derived from the WT and Y86* strains. The peak at 16.198 min corresponding to m^1^G is indicated by the dotted line.

TrmD is a high-priority antibiotic target and development of TrmD inhibitors is currently an active area of research (15–21). Several factors have established TrmD as a promising antibiotic target. Firstly, the *trmD* gene is widely conserved across bacteria and has been reported to be indispensable for growth in phylogenetically diverse species including *E. coli* (11), *Streptococcus pneumoniae* (22), *Pseudomonas aeruginosa* (10) and *Mycobacterium abscessus* (17). Secondly, there is no homolog of TrmD in humans; although the m^1^G37 modification is also conserved in archaea and eukaryotes, it is introduced by the unrelated Rossmann-fold methyltransferase TRM5, which is phylogenetically, structurally, and mechanistically distinct from TrmD (5). Thirdly, knockdown of TrmD has been shown to have widespread, pleiotropic effects on cell viability, which could suppress the evolution of antibiotic resistance (13, 21). For example, frameshift errors induced by loss of m^1^G37 are known to have a disproportionate impact on membrane proteins, since slowly-translated and frameshift-prone CC[C/U]-[C/U] motifs occur frequently near the start of sequences encoding membrane proteins, perhaps as a mechanism to promote insertion of the nascent polypeptide into the membrane (13). As a result, loss of m^1^G37 impairs expression of membrane proteins, which disrupts membrane structure, increases membrane permeability, impairs membrane efflux, and increases susceptibility to various antibiotics (13). Based on present knowledge, it is unclear how bacteria could evolve to tolerate such widespread impacts on cell physiology.

Although it has been established beyond doubt that the m^1^G37 modification improves translational accuracy, it is less clear that frameshift errors induced by loss of m^1^G37 are responsible for the essentiality of TrmD in bacteria. Several considerations hinted that TrmD might have a broader role than prevention of translational errors and might be dispensable for growth under certain conditions. Firstly, m^1^G37 is known to have a broader role than preventing frameshift errors in archaea and eukaryotes; for example, it is required for recognition of yeast tRNA^Asp^ (GUC) and *Methanocaldococcus jannaschii* tRNA^Cys^ by their respective aminoacyl-tRNA synthetases (23, 24). Secondly, there is some limited evidence that bacteria have a high level of tolerance for frameshift errors. While estimates for the basal rate of translational errors in bacteria vary depending on the species, reporter system, codon context, and growth conditions, in some cases frameshift error rates as high as ~2–10% per codon have been observed (25, 26). Although loss of m^1^G37 in tRNA^Pro^ can increase the frameshift error rate at Pro codons up to eight-fold in the worst-case scenario (specifically, at a CCC-C sequence at the second position of an open reading frame), in most contexts, the effect of m^1^G37 loss is fairly small in magnitude (less than five-fold) (11, 27); it is therefore surprising that frameshift errors induced by loss of m^1^G37 result in a complete loss of cell viability. Finally, some observations indicate that there may be mechanisms that enable bacteria to survive in the absence of TrmD. For example, *E. coli* strains with the G37C mutation in tRNA^Pro^ (UGG) show slow growth even though the mutant tRNA lacks m^1^G37 (13). Furthermore, *TRM5* in yeast was identified in a genome-wide screen as an essential gene whose essentiality can be overcome by adaptive evolution (28); this might also be true of the analogous gene *trmD* in bacteria. Based on these considerations, we hypothesized that the prevention of frameshift errors might not be the main cause of growth defects observed in the absence of m^1^G37 and that it might be possible for bacteria to evolve rapidly in response to loss of m^1^G37.

Evolutionary repair offers a methodology to address both of these hypotheses. In evolutionary repair experiments, a microbial strain is subjected to a genetic perturbation, usually gene deletion, and then propagated over hundreds to thousands of generations to allow the population to evolve in response to the perturbation (29). Next-generation sequencing (NGS) of population genomic DNA or whole-genome sequencing of isolated clones can be used to identify any adaptive mutations that alleviate the fitness defect caused by the original perturbation. However, application of this methodology to essential genes presents an obvious challenge, which is that essential genes by definition cannot be deleted. Rodrigues and Shakhnovich recently presented a useful solution to this problem, whereby chromosomal point mutations are introduced to disrupt but not completely inactivate an essential protein (30). The choice of mutation at the DNA level dictates which amino acids are accessible by a single nucleotide mutation, which can be used to control the ease of reversion of the engineered mutation in the evolutionary experiment (30). Although evolutionary repair is conceptually similar to the classical genetic method of suppressor analysis, which aims to identify individual mutations that compensate for the deleterious effect of a mutation in a gene of interest, evolutionary repair experiments provide a much more detailed view of the cumulative effects of multiple mutations, the relative fitness effects of different mutations, and the dynamics of the evolutionary process (29).

In this work, we applied this experimental evolution approach to understand whether *E. coli* can adapt in response to m^1^G37 deficiency, in order to better understand the biological function of this essential tRNA modification. We engineered seven *E. coli* strains with different mutations in the genomic copy of *trmD*. Surprisingly, strains bearing substitutions in catalytic residues and premature stop codons were viable despite the absence of m^1^G. Experimental evolution of five mutant strains in a total of 15 independent populations showed that *E. coli* recovers rapidly and reproducibly in response to m^1^G37 deficiency. NGS of population genomic DNA revealed that mutations in the proline-tRNA ligase gene *proS* were mainly responsible for recovery of strains that showed a complete loss of TrmD activity. Together with growth assays and *in vitro* aminoacylation assays, these results showed that m^1^G37 affects the aminoacylation rate of tRNA^Pro^ by the ProRS enzyme and that growth defects associated with m^1^G37 deficiency can be largely countered by point mutations in *proS* that improve aminoacylation of unmodified tRNA.

## Results

### Construction and analysis of *trmD* mutant strains

In order to perform experimental evolution of *trmD*-deficient *E. coli* strains, our first goal was to construct *trmD* mutant strains that showed severe growth defects relative to wild-type *E. coli* but were still viable. Based on literature data on the growth of bacteria with mutations in *trmD* and catalytic activity of TrmD variants *in vitro* (summarized in Supplementary Table 1), we chose six substitutions to introduce into the genomic copy of *trmD*. These included two substitutions in key catalytic residues (R154A and D169A), which largely abolish catalytic activity, two substitutions in active site residues (Y115A and G141A), which have more modest effects on catalytic activity, and two substitutions (G117N and S165L) that have been reported to reduce colony size in *Salmonella enterica* by at least 90% (3, 7, 31, 32). At the DNA level, we generally designed the mutations such that two independent nucleotide mutations would be required for reversion to the wild-type amino acid (Supplementary Table 2). In some cases, a viable mutational trajectory to the wild-type amino acid could be predicted. For example, the substitution D169A can effectively be reverted by a single nucleotide mutation yielding Glu, which gives enzyme activity equivalent to the wild-type (WT) Asp variant (31, 32). On the other hand, in the case of R154A, no single nucleotide mutations were anticipated to restore enzyme activity.

As a negative control, we also constructed a translational knockout strain of *trmD* (Y86*), in which three consecutive codons from Tyr86 were replaced with three different stop codons to prevent functional TrmD expression while avoiding readthrough of the stop codons *via* nonsense suppressor mutations (33). Compared with traditional methods for gene deletion, the translational knockout strategy has the advantage of minimizing disruption of neighboring genes, which is particularly important in this case because *trmD* is co-transcribed with the essential and highly expressed ribosomal genes *rpsP* and *rplS* (34). Indeed, some *trmD* deletion mutants have shown altered expression of RplS (34), and previous studies have shown that gene deletion can cause unexpected growth defects and compensatory mutations in evolutionary experiments, e.g., by disrupting translation of neighboring genes (35).

The seven genomic mutations were introduced into *E. coli* BW25113 cells complemented with an arabinose-inducible yeast TRM5 expression vector by recombineering. Each engineered strain contained a mutation in *trmD* and a unique barcode sequence downstream of *glnH* so that the evolved strain could be distinguished easily from WT *E. coli* even if the mutation in *trmD* was reverted. After genome mutagenesis, the yeast TRM5 expression vector was removed using a CRISPR/Cas9-based plasmid curing system (36). Plasmid-cured colonies of all mutant strains appeared on Luria-Bertani (LB) medium agar plates within 36 h at 37 °C.

The effects of the *trmD* mutations were compared by growth assays of the mutant strains in liquid culture (LB medium, 37 °C; Fig. 1b). The Y115A and G141A substitutions had little effect on growth. The R154A, S165L, D169A, and Y86* mutants showed a severe reduction in growth rate compared with the WT strain (2.8-fold, on average), while the G117N substitution had an intermediate effect (1.7-fold) (Supplementary Fig. 1). Surprisingly, in contrast to previous reports of the essentiality of *trmD* in *E. coli* (11, 37), both the translational knockout strain (Y86*) and catalytic mutant strains (R154A and D169A) were viable. To confirm that the viability of these strains was not a result of residual TrmD activity, we performed UPLC analysis of nucleoside content in tRNA^Pro^ (CGG) purified from the strains that showed growth defects. m^1^G was detected in tRNA purified from the G117N, S165L, and D169A strains (<10% of WT), but not the R154A or Y86* strains (below detection limit, <1% of WT) (Fig. 1c–d, Supplementary Fig. 2). These results indicate that although *E. coli* strains lacking m^1^G exhibit very slow growth, m^1^G is not completely indispensable for cell viability, and that the apparent essentiality of *trmD* might be an artefact of recombination-based methods for construction of *trmD* knockout strains.

Next, we performed experimental evolution of the five strains that showed growth defects. For each strain, three separate populations initialized from different colonies were grown in batch culture with daily serial passaging for 15 to 30 days. Samples were taken periodically for glycerol stock storage and extraction of genomic DNA, and growth assays were performed weekly to monitor recovery of growth. The G117N and D169A mutant populations showed rapid recovery, returning to a growth rate equivalent to the WT strain within one week (~47 generations), while the R154A, S165L, and Y86* mutant populations showed a rapid initial recovery within one week followed by slower recovery over the next several weeks, although in general they had not attained the growth rate of the WT strain by the end of the experiment (range of growth rates for final R154A, S165L, and Y86* populations: 1.31 to 1.57 h^−1^, versus 1.53 h^−1^ for WT) (Fig. 2, Supplementary Fig. 1). Overall, these results show that *E. coli* adapts rapidly and reproducibly to deficiency of m^1^G37 caused by mutagenesis of the *trmD* gene.

**Figure 2.**
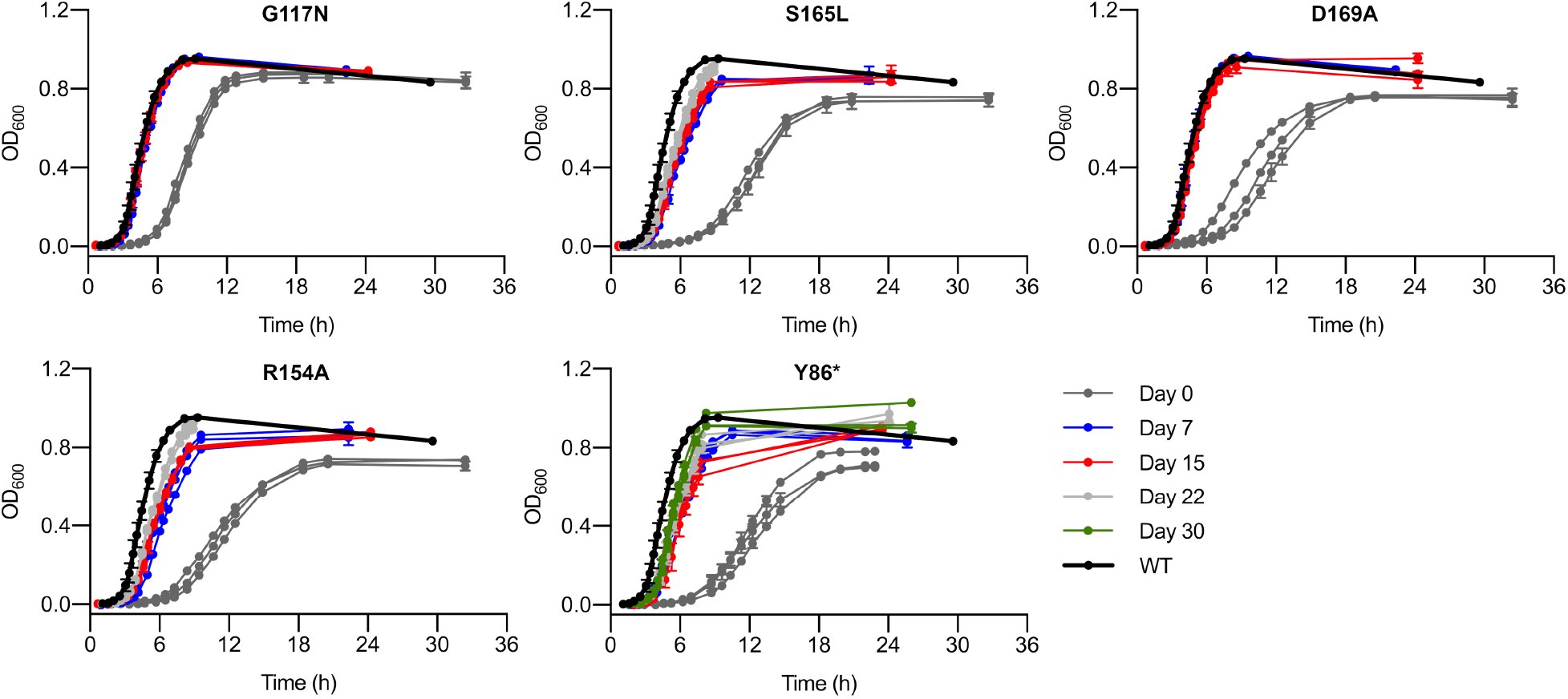
Experimental evolution of *trmD* mutant strains. Growth curves of each population taken at the indicated time points of the evolutionary experiment are shown. Three replicate populations are shown for each strain. Data represent mean ± s.d. of two technical replicates for each population. Data for day 0 are reproduced from Fig. 1b for comparison.

### Mechanisms for adaptation towards *trmD* deficiency

We next aimed to identify the genomic changes underlying adaptation to *trmD* deficiency. Firstly, we attempted to eliminate the possibility that recovery had occurred by trivial mechanisms, such as simple reversion of the engineered *trmD* mutations. Sanger sequencing of the portion of the *trmD* gene containing the engineered mutations in the evolving populations showed that the D169A populations all recovered *via* the substitution A169E, which is known to restore TrmD activity (31, 32); thus, these populations were not analyzed further. Sanger sequencing also revealed the presence of coding mutations in *trmD* in some of the G117N and S165L populations (discussed further below), but not in the R154A and Y86* populations. We also suspected that tandem amplification of *trmD* might have occurred as another trivial mechanism for recovery in strains with point mutations in *trmD*. We hypothesized that amplification of *trmD* may have amplified the residual enzyme activity by increasing gene dosage and enzyme expression, explaining the rapid increases in fitness observed in the evolutionary experiment. Indeed, gene duplication occurs orders-of-magnitude more frequently than beneficial point mutations in a given gene (roughly 10^−5^ to 10^−2^ and 10^−9^, respectively) (38), and evolutionary experiments have shown that gene amplification can drive rapid increases in fitness when a low-level enzyme activity is rate-limiting for growth (35). However, measurement of the average copy number of *trmD* in population genomic DNA by quantitative PCR (qPCR) at five different time points (days 1, 3, 6, 15 and 22) showed that the gene remained at a single copy in the G117N, S165L, D169A, and R154A populations throughout the evolutionary experiment (Supplementary Fig. 3).

We next performed NGS of population genomic DNA isolated from each population at the end of the evolutionary experiment (day 15, 22 or 30), and except for the G117N populations, from an early intermediate population (day 9). The mutations observed in each population are summarized in Figure 3a, and complete lists of mutations and genomic rearrangements are given in Supplementary Tables 3 and 4, respectively. Firstly, some mutations were observed with 100% frequency in all populations derived from a single mutant strain (7–9 per strain), which were inferred to represent off-target mutations introduced during strain construction. The presence of these mutations in the original mutant strains was confirmed by targeted Sanger sequencing of the genomic region surrounding the mutation. The mutant strains showed growth similar to the WT strain when complemented with the yeast TRM5 expression vector, indicating that the off-target mutations were not responsible for the decreased growth rate of the *trmD* mutant strains and were therefore unlikely to impact adaptation to m^1^G37 deficiency (Supplementary Fig. 4). The fact that the same adaptive mutations were frequently observed in different *trmD* mutant strains with different off-target mutations is another indication that the off-target mutations had no significant effect on the evolutionary outcome.

**Figure 3.**
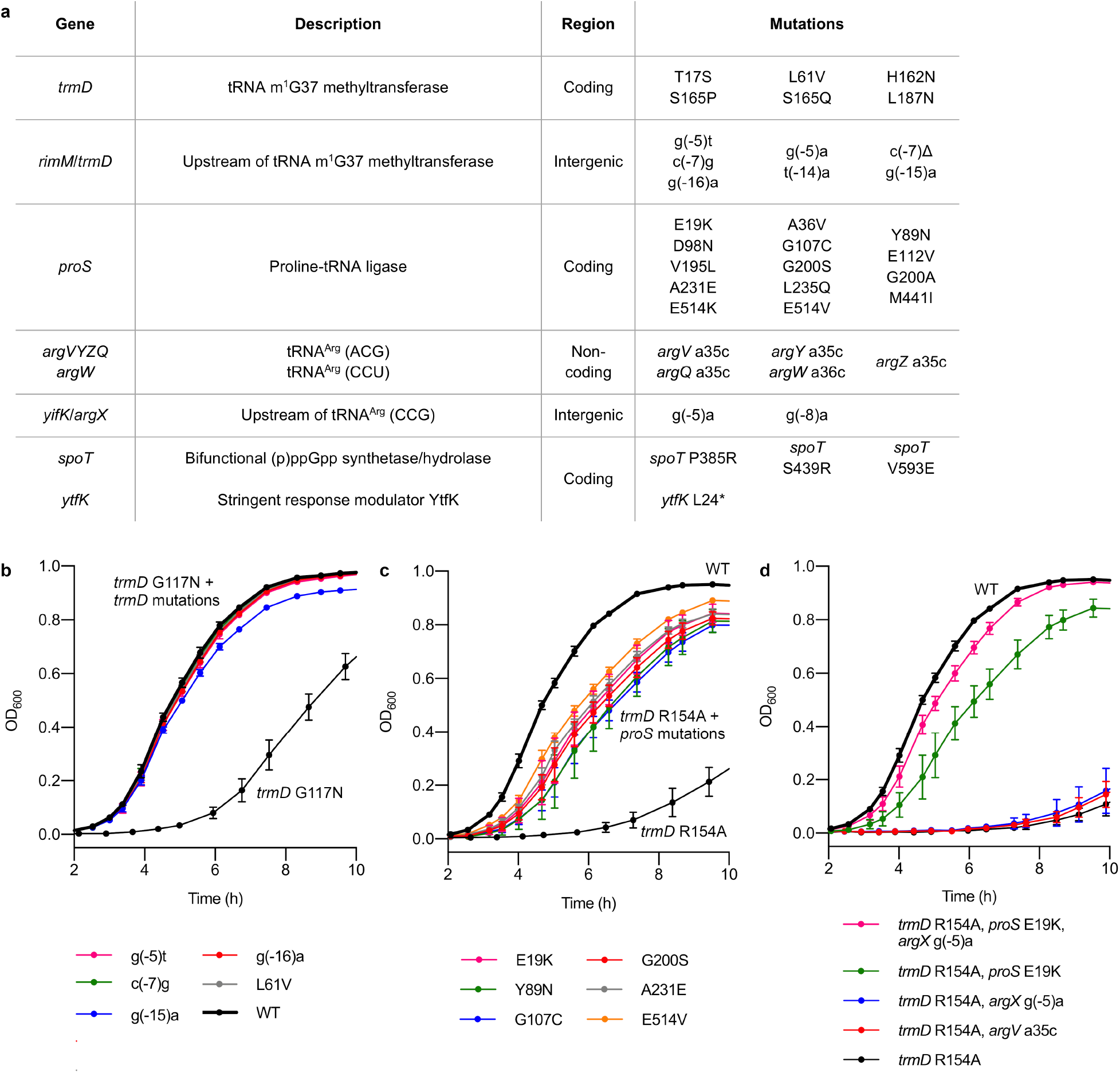
Mechanisms of adaptation to *trmD* deficiency. **a,** Summary of the six main categories of mutations observed in the evolutionary experiment. **b-d,**Growth assays of single and double mutants of original *trmD* mutant strains in LB media at 37 °C, showing that single mutations in *trmD* and *proS* can substantially improve growth of the *trmD* G117N and R154A strains, respectively. Data represent mean ± s.d. for the indicated number of biological replicates (different colonies). **b,** Variants of the *trmD* G117N strain with an additional mutation in *trmD*. WT, g(−5)t, c(−7)g, g(−15)a, *n* = 3; g(−16)a, L61V, *n* = 6 (three colonies each from two independently isolated strains). Data for *trmD* G117N are reproduced from Fig. 1b for comparison. **c,** Variants of the *trmD* R154A strain with an additional mutation in *proS*. WT, E514V, *n* = 3; G200S, A231E, E19K, Y89N, G107C, *n* = 6 (three colonies each from two independently isolated strains). Additional replicates of this experiment are shown in Supplementary Figure 5. Data for the *trmD* R154A mutant is reproduced from Fig. 1b for comparison. **d,** Variants of the *trmD* R154A strain with mutations in *argX* and *argV*. WT, *trmD* R154A, *n* = 3; other strains, *n* = 6 (three colonies each from two independently isolated strains). Data for the WT and *trmD* R154A, *proS* E19K strains are duplicated from Fig. 3b because the triple mutant was grown on the same plate. The three remaining mutants were assayed in a separate experiment.

In summary, the main categories of adaptive mutations observed across the evolving populations were: (1) coding mutations in *trmD*; (2) non-coding mutations upstream of *trmD*; (3) coding mutations in *proS*, which encodes the proline-tRNA ligase ProRS and is responsible for aminoacylation of tRNA^Pro^ for use in protein translation; (4) non-coding mutations in the tRNA^Arg^ genes *argVYZQ* and *argW*; (5) non-coding mutations upstream of the tRNA^Arg^ gene *argX*; and (6) coding mutations in genes associated with the guanosine (penta-)tetraphosphate ((p)ppGpp)-mediated stringent response. The final G117N populations showed three coding mutations in *trmD* and three non-coding mutations upstream of *trmD*. One population showed fixation of a single mutation, *trmD* g(−15)a, while the other two populations showed a mixture of mutations with a total frequency of >85%. The final R154A and Y86* populations showed no mutations in *trmD*, as expected based on the Sanger sequencing data; instead, mutations in *proS* were observed with a high frequency. In two populations, fixation of a single amino acid substitution (A231E) was observed, while the other populations showed a mixture of ProRS substitutions with a total frequency of >85%. Mutations in *argQZYV* or *argW*, or upstream of *argX*, were also observed in all populations, with total frequencies varying from 9% to 80%. Missense mutations in *spoT* and a nonsense mutation in *ytfK*, which are both associated with the stringent response, were observed in some populations with a frequency between 9% and 57%. Finally, the final S165L populations showed a combination of the mutations observed in the G117N, R154A, and Y86* populations. All populations showed mutations upstream of *trmD* (total frequency 22% to 85%). In addition, one population showed coding mutations in *trmD*, one population showed a coding mutation in *proS*, and one population showed mutations in *proS* and *argV*. Altogether, these results show that the G117N and S165L strains, which have some residual TrmD activity (Fig. 1c), can recover through compensatory mutations in *trmD*, while the R154A and Y86* strains, which show severe and irreversible defects in TrmD (substitution of a catalytic residue and premature stop codons, respectively), recover primarily through mutations in *proS*.

To investigate the adaptive role of individual mutations in more detail, we obtained representative double mutant strains containing a single mutation in addition to the original *trmD* mutation, either by isolating individual strains from the evolved populations or by introducing genomic mutations into the *trmD* mutant strains by recombineering. Growth analysis of five variants of the *trmD* G117N strain containing a second mutation in *trmD* (g(−16)a, g(−15)a, c(−7)g, g(−5)t or L61V) confirmed that a single point mutation was sufficient for complete recovery of growth of the G117N mutant (Fig. 3b). Likewise, isolation of six variants of the *trmD* R154A strain containing a single mutation in *proS* (E19K, Y89N, G107C, G200S, A231E, or E514V) showed that a single mutation in *proS* is sufficient to substantially recover the growth of the *trmD* R154A strain (range of growth rates of *proS* mutants: 1.11 to 1.24 h^−1^, compared with 1.48 h^−1^ for WT and 0.60 h^−1^ for *trmD* R154A) (Fig. 3c, Supplementary Figs. 1 and 5). On the other hand, the *argX* g(− 5)a and *argV* a35c mutations had no effect on growth of the *trmD* R154A mutant, but the *argX* g(−5)a mutation did improve growth of the *trmD* R154A/*proS* E19K double mutant strain (Fig. 3d). This observation is consistent with the fact that mutations in tRNA^Arg^ genes were only observed in populations that also contained *proS* mutations at a high frequency. Altogether, these results provide further evidence that the mutations in *trmD*, *proS*, and *argX* observed in the evolutionary experiment occur as an adaptive response to m^1^G37 deficiency and suggest that mutation of *proS* is the dominant mechanism for growth recovery in mutant strains where TrmD activity cannot be recovered through single point mutations.

### Evolutionary dynamics of the response to m1G37 deficiency

Next, we analyzed the dynamics of the evolutionary response to m^1^G37 deficiency by comparison of the mutations observed in the intermediate populations (day 9) and the final populations (Fig. 4a). To gain further qualitative insight into the early stages of the evolutionary trajectory, we also performed Sanger sequencing of the coding region of *proS* in population genomic DNA samples from days 1, 3, and 6 to identify any polymorphisms (Supplementary Fig. 6). Together, these results showed that the early evolutionary trajectory is dominated by competition between strains with different mutations in *proS*. Fourteen different substitutions in ProRS were observed, with six arising independently in multiple populations or affecting the same protein residue, which suggests more strongly that they have an adaptive effect. The S165L, R154A, and Y86* intermediate populations show between one and four different mutations in *proS*, with a total frequency of >75% (except the Y86*-3 population, which instead shows duplication of *proS*; see below). Between the intermediate and final populations, competition between strains with different mutations in *proS* causes eventual dominance of a single substitution, with E19K, A231E, E514K, and E514V showing high frequencies at the end of the evolutionary experiment, while substitutions such as Y89N and E112V are outcompeted. Although the persistence of these substitutions could be explained by genetic hitchhiking rather than high fitness, the E19K, A231E, and E514V substitutions also show a slight fitness advantage over the G107C substitution in growth assays (Fig. 3c, Supplementary Figs. 1 and 5). Altogether, these results indicate that *proS* is a large mutational target in the evolutionary response to m^1^G37 deficiency, that is, there are many different amino acid substitutions in ProRS that can provide an initial pathway to recovery before they are outcompeted by substitutions with higher fitness.

**Figure 4.**
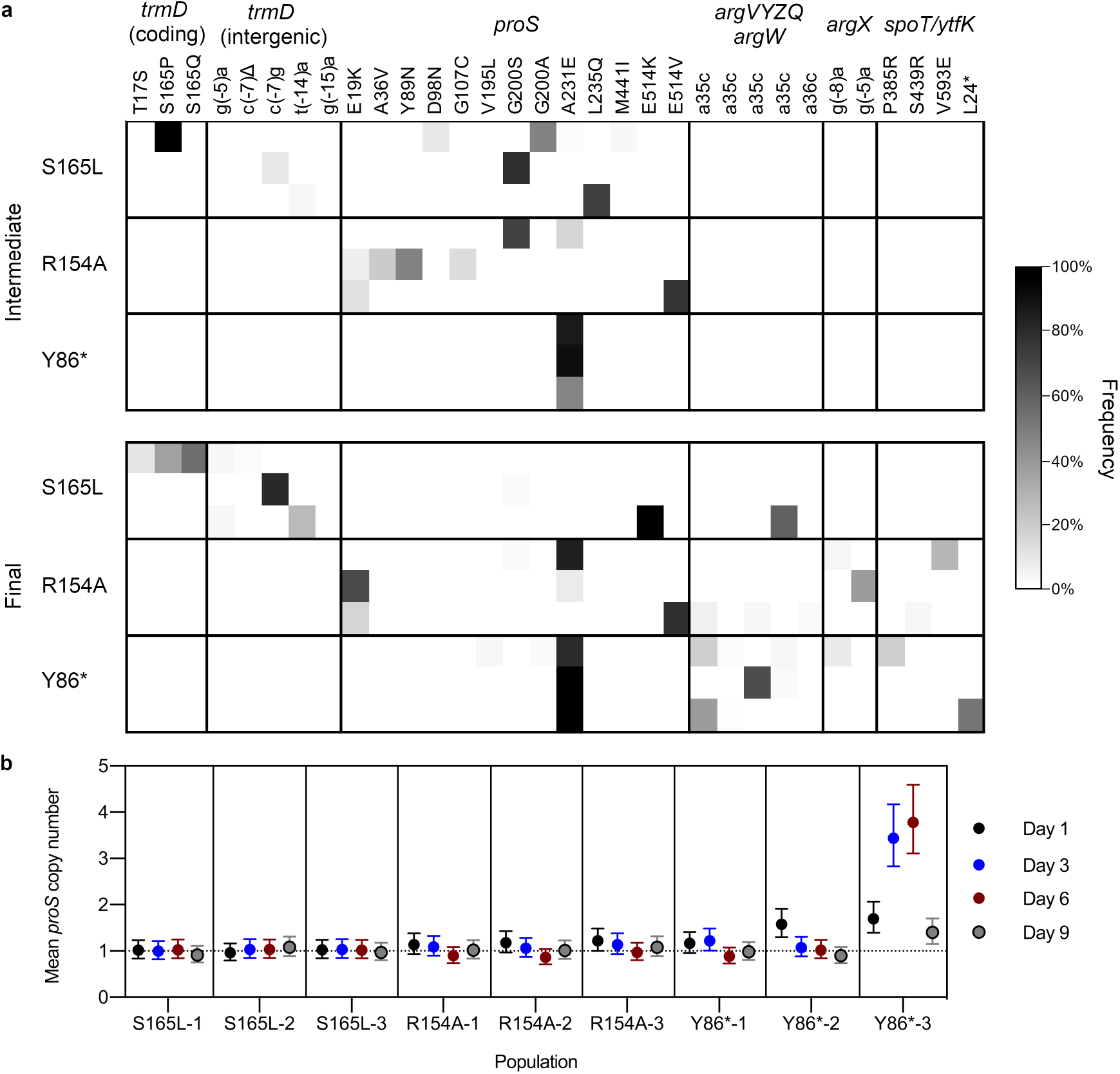
Evolutionary dynamics of adaptation to *trmD* deficiency. **a,** Comparison of mutation frequencies in each S165L, R154A, and Y86* population at an intermediate (day 9) and final time point (day 22 or day 30). Each row represents one population, each column represents one mutation, and the shading represents the observed frequency of the mutation. Mutations observed only in the G117N populations are not shown. **b,** Mean copy number of *proS* in population genomic DNA from the S165L, R154A, and Y86* populations at different time points, measured by qPCR. Error bars represent 95% confidence intervals. Estimates are derived from three technical replicates per primer pair for each sample. Significance testing was performed by one-way ANOVA with Dunnett’s test for multiple comparisons using a signifiance threshold of *P* < 0.05. The following populations showed a statistically significant increase in *proS* copy number compared with WT *E. coli*: R154A-3 (day 1) (*P* = 0.0400), Y86*-1 (day 3) (*P* = 0.0371), Y86*-2 (day 1) (*P* < 0.0001), and all Y86*-3 time points (*P* < 0.0001).

Comparison of the mutations observed in the intermediate and final populations also shows a major difference in the trajectories of the S165L populations and the R154A and Y86* populations. Between the intermediate and final R154A and Y86* populations, mutations in *proS* are retained, and further mutations occur in tRNA^Arg^ genes and genes associated with the stringent response. However, in two of the final S165L populations, mutations in *proS* are purged from the population and are replaced with mutations in *trmD*. Mutations in *trmD* are present at a low frequency (<15% in total) in the intermediate S165L populations, indicating that competition between strains bearing different mutations, rather than reversion of *proS* mutations after fixation of *trmD* mutations, is the most likely explanation for the decrease in frequency of *proS* mutations. This result provides another indication that the dynamics of the evolutionary response to m^1^G37 deficiency is dominated by the large mutational target size of *proS*; because there are more mutations that compensate for the effect of the S165L substitution in *proS* than in *trmD*, they appear with a high frequency at the early stages of the evolutionary trajectory even though they provide a smaller fitness benefit. Altogether, these results show that initial adaptation of *E. coli* to m^1^G37 deficiency occurs by mutations in *proS*. If there is a viable pathway to increase TrmD activity through coding or promoter mutations, then these mutations outcompete the mutations in *proS*; otherwise, mutations in *proS* are retained and further gains in fitness are realized by mutations in tRNA^Arg^ genes and genes associated with the stringent response.

In addition to coding mutations in *proS*, we also observed insertion-sequence-mediated duplication of an 82.3 kb fragment containing *proS* in one population (Y86*-3-9), with a frequency of 28%. The boundaries of the duplication were identified by analysis of the new junction sequences and confirmed by analysis of read coverage depth (Supplementary Fig. 7). To study the dynamics of gene amplification early in the evolutionary trajectory and to determine whether amplification of *proS* also occurred in other populations, we also quantified the mean copy number of *proS* in population genomic DNA of the S165L, R154A and Y86* populations at early time points (days 1, 3, 6, and 9) by qPCR (Fig. 4b). Consistent with the NGS data, only the Y86*-3 population showed a significant increase in *proS* copy number at day 9 (1.40, *P* < 0.0001 by one-way ANOVA with Dunnett’s test for multiple comparisons). Earlier in the evolutionary trajectory, a substantial increase in *proS* copy number was also observed in population Y86*-2 at day 1 (1.58, *P* < 0.0001). In population Y86*-3, rapid amplification and deamplification of *proS* was observed, with mean copy number reaching a maximum of 3.78 at day 6 before decreasing rapidly to 1.40 at day 9. Notably, polymorphisms in *proS* are not observed in the Y86*-3 population until day 9 (Supplementary Fig. 6), indicating that amplification of *proS* precedes mutation and deamplification.

The fact that amplification of *proS* is followed by mutation and deamplification indicates that the evolution of *proS* in the Y86*-3 population proceeds by an innovation-amplification-divergence (IAD) mechanism (38, 39). IAD refers to an evolutionary mechanism that occurs when a low-level secondary function of a gene (for example, a promiscuous enzyme activity) becomes important for organismal fitness. In this scenario, gene amplification can provide an immediate selective benefit by amplifying the secondary function of the gene. Incidentally, amplification of a gene increases the probability of mutations occurring in that gene, accelerating the fixation of adaptive mutations that improve the secondary function. After these adaptive mutations occur, selective pressure for gene amplification is relaxed and the extra gene copies are lost. In this case, the observation that *proS* can evolve by an IAD mechanism has two main implications: firstly, it suggests that the fitness benefits of the coding mutations observed in *proS* are associated with an increase in a low-level enzyme activity, rather than, for example, a decrease in aminoacylation activity to activate the stringent response, a common adaptive mechanism in antibiotic tolerance (40). Aminoacylation of non-native G37-unmodified tRNA is a likely candidate for the low-level enzyme activity of ProRS that is amplified *via* the IAD mechanism. Secondly, it suggests another reason (in addition to the large mutational target size of *proS*) that coding mutations in *proS* can occur very rapidly in response to m^1^G37 deficiency. It should be noted that amplification of large fragments of genomic DNA carries a significant fitness cost (38, 41) and it is therefore implausible that amplification of *proS* to a copy number of ~4 would occur if gene amplification did not provide a significant fitness benefit.

### Adaptation to *trmD* deficiency *via* mutations in *proS*

Because mutations in *proS* were the dominant mechanism for recovery of strains in which TrmD activity cannot be recovered, we attempted to investigate the effects of these mutations at the biochemical level. The simplest explanation for the observation that amplification and mutation of *proS* improve growth of *trmD* mutant strains is that ProRS is unable to efficiently aminoacylate tRNA^Pro^ lacking m^1^G37, and that substitutions in ProRS increase the aminoacylation efficiency of G37-unmodified tRNA^Pro^. Although previous work has suggested by indirect reasoning that ProRS does not distinguish between m^1^G37-modified and G37-unmodified tRNA^Pro^ (42, 43), the effect of the m^1^G37 modification in tRNA^Pro^ on the aminoacylation activity of ProRS has not been measured directly. We therefore performed *in vitro* aminoacylation assays of purified ProRS variants with m^1^G37-modified and unmodified tRNA^Pro^ (CGG), which is the tRNA^Pro^ species with the highest concentration in *E. coli*. We used tRNA^Pro^ (CGG) purified from wild-type *E. coli* or an evolved *trmD* mutant strain, ensuring that the tRNA was fully modified except at position 37 and allowing us to isolate the effect of the m^1^G37 modification. Three ProRS variants with substitutions in different parts of the protein structure (E19K, G200S and E514V) were selected for characterization (Fig. 5a).

**Figure 5.**
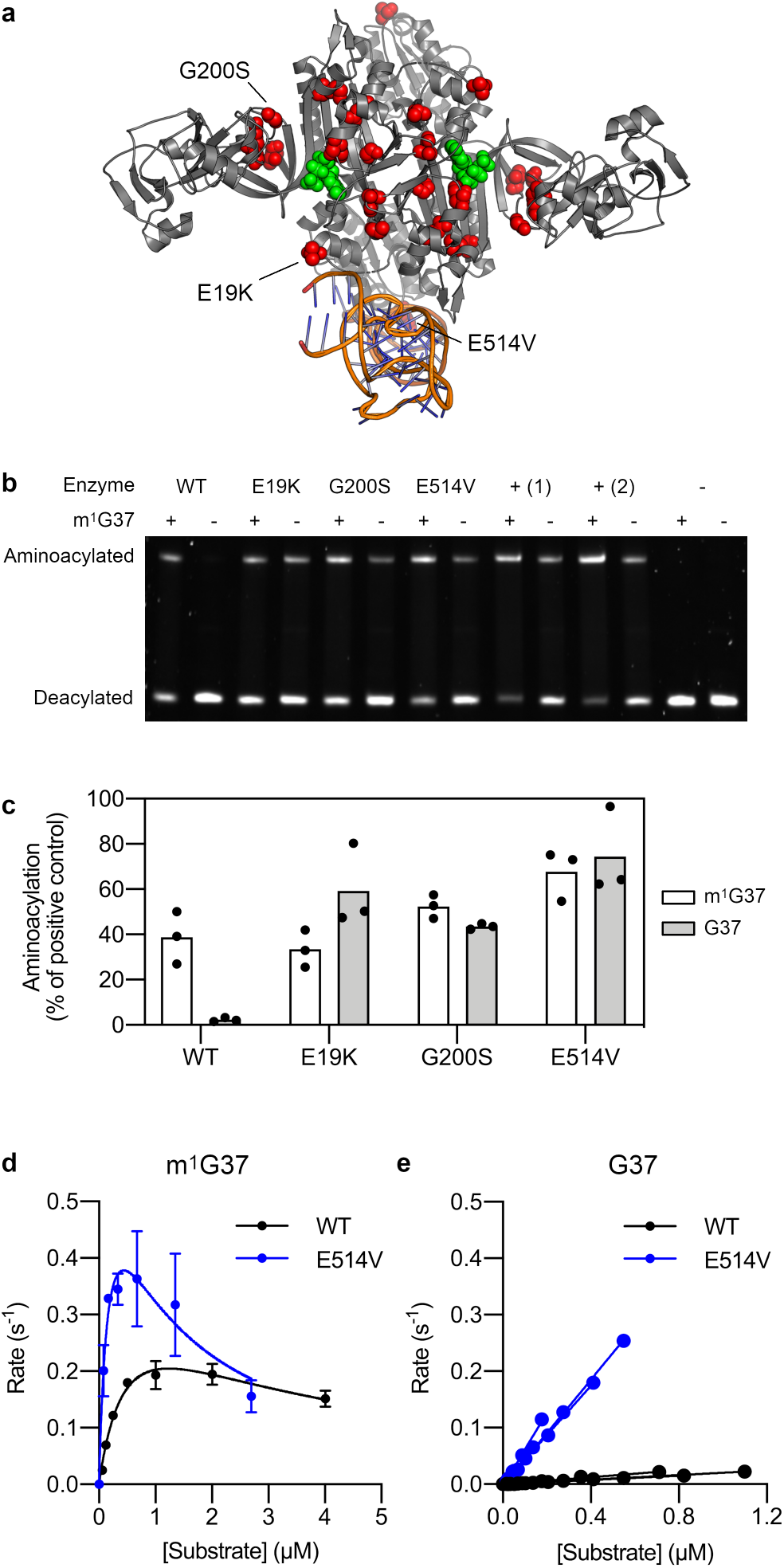
Mutations in *proS* allow efficient aminoacylation of tRNA^Pro^ lacking m^1^G37. **a,** Structural positions of ProRS substitutions observed in the evolutionary experiment. Positions where substitutions were observed are shown as red spheres in the structure of ProRS from *Pseudomonas aeruginosa* (71% identity to *E. coli* ProRS, PDB: 5UCM). The tRNA and ATP (green spheres) structures are derived from the tRNA-bound structure of ProRS from *Thermus thermophilus* (16% identity to *E. coli* ProRS, PDB: 1H4Q); this structure was superimposed onto the *P. aeruginosa* structure to show the approximate positions of the tRNA binding site and active site. **b-c,** Comparison of aminoacylation activity of ProRS variants on m^1^G37-modified and G37-unmodified tRNA^Pro^ (CGG) using a gel shift assay. Reaction conditions: tRNA concentration 2 μM, enzyme concentration 2.5 nM, reaction temperature 37 °C, reaction time 14 min. The enzyme concentration was chosen to ensure linearity of product formation for the WT enzyme with m^1^G37-modified tRNA over the course of the assay. **b,** + (1), positive control with 1 μM WT ProRS; + (2), positive control with 1 μM ProRS E514V; −, no enzyme control. This gel image was taken with a high exposure to increase visibility of the bands; a lower exposure was used for data analysis. Annotated and original raw gel images are provided in Supplementary Fig. 8 and Supplementary Data 1, respectively. **c,**Mean and individual values of three replicates from separate experiments. Significance testing was performed by one-way ANOVA with Dunnett’s test for multiple comparisons using a significance threshold of *P* < 0.05. WT m^1^G37 vs. WT G37, *P* = 0.0073; WT m^1^G37 vs. E19K m^1^G37, *P* = 0.9897 (n.s.); WT m^1^G37 vs. G200S m^1^G37, *P* = 0.5622 (n.s.); WT m^1^G37 vs. E514V m^1^G37, *P* = 0.0371. **d-e**, Michaelis-Menten kinetic analysis of tRNA^Pro^ (CGG) aminoacylation by the WT and E514V ProRS variants. Representative annotated and original raw gel images are provided in Supplementary Fig. 9 and Supplementary Data 1, respectively. Kinetic parameters are summarized in Supplementary Table 5. **d,** m^1^G37-modified tRNA. Data represent mean ± s.d. of two technical replicates from a single representative experiment. **e,**G37-unmodified tRNA. Results from three separate experiments are shown for each variant.

Firstly, we performed a direct comparison of aminoacylation efficiency between the ProRS variants using an end-point assay at a fixed tRNA concentration (2 μM) within the physiological concentration range (44) (Fig. 5b–c). Consistent with our hypothesis, replacement of m^1^G37-modified tRNA with G37-unmodified tRNA resulted in a 17-fold reduction in aminoacylation in the case of the WT enzyme. In contrast, the E19K, G200S, and E514V variants showed similar or greater aminoacylation efficiency on m^1^G37-modified tRNA compared with the WT enzyme (0.9− to 1.7-fold difference, variant to WT ratio), while also showing a similar aminoacylation efficiency between m^1^G37-modified tRNA and G37-unmodified tRNA (0.6− to 1.2-fold difference, m^1^G37 to G37 ratio). Consistent with the observation that substitutions in ProRS had no effect on the aminoacylation efficiency of m^1^G37-modified tRNA, we found that *E. coli* strains bearing the same mutations in the genomic copy of *proS* showed growth indistinguishable from the WT strain (Supplementary Fig. 10).

To gain more quantitative insight into the differences in aminoacylation efficiency, we also performed Michaelis-Menten kinetic analysis of the WT and E514V variants, although for G37-unmodified tRNA, only *k*_cat_/*K*_M_ could be determined due to the low tRNA yield and high *K*_M_ values (Fig. 5d–e, Supplementary Table 5). As expected, the WT enzyme showed a severe reduction in catalytic efficiency for G37-unmodified tRNA compared with m^1^G37-modified tRNA (17.5-fold reduction in *k*_cat_/*K*_M_). The E514V variant showed higher catalytic efficiency than the WT enzyme on both m^1^G37-modified tRNA (4.5-fold increase in *k*_cat_/*K*_M_) and G37-unmodified tRNA (17.2-fold increase in *k*_cat_/*K*_M_); in the case of m^1^G37-modified tRNA, this was a result of both a decrease in *K*_M_ (0.54 μM to 0.27 μM) and an increase in *k*cat (0.40 to 0.84 s^−1^). The aminoacylation efficiency of the E514V variant on G37-unmodified tRNA was very similar to the aminoacylation efficiency of the WT enzyme on m^1^G37-modified tRNA (*k*_cat_/*K*_M_ 5.22 × 10^5^ M^−1^ s^−1^ vs. 7.31 × 10^5^ M^−1^ s^−1^). Altogether, these results show that loss of the m^1^G37 modification dramatically impairs aminoacylation of tRNA^Pro^ by ProRS, but that aminoacylation of unmodified tRNA can be restored by a single substitution in ProRS.

### Adaptation to *trmD* deficiency *via* other mechanisms

Although we focused on biochemical characterization of mutations in *proS* because these represented the main mechanism for recovery of TrmD-deficient strains, the mutations observed elsewhere in the genome also provide insight into m^1^G37 function, *trmD* regulation, and the evolutionary response to m^1^G37 deficiency. In the case of the G117N and S165L populations, coding mutations in *trmD* presumably compensate for the deleterious effects of the engineered G117N and S165L substitutions, while mutations upstream of *trmD* presumably increase enzyme expression to compensate for decreased catalytic activity. It is known that TrmD is expressed at a much lower level (~40-fold lower) than RpsP and RplS, which are co-transcribed with *trmD*; this mechanism enables TrmD, RpsP and RplS regulation to be coordinated while allowing the ribosomal proteins RpsP and RplS to be expressed at a higher level (45). The disparity in expression levels can be explained by unusually strong mRNA secondary structure at the 5’ end of the *trmD* ORF that inhibits translation (46–48). The mutations upstream of *trmD* observed in the evolutionary experiment are predicted to disrupt this mRNA secondary structure, increasing translation (Supplementary Fig. 11). Thus, the unusual mechanism for translational regulation of *trmD* appears to be responsible for the existence of a large number of mutations that can increase TrmD expression.

In the S165L, R154A, and Y86* populations, mutations in tRNA^Arg^ genes and genes involved in the stringent response frequently appear after mutations in *proS*. *E. coli* has seven tRNA^Arg^ genes, of which only tRNA^Arg^ (CCG) (*argX*) contains the m^1^G37 modification. The four tRNA^Arg^ (ACG) genes (*argVYZQ*) have the same anticodon sequence and are redundant. tRNA^Arg^ (CCG) (*argX*), tRNA^Arg^ (CCU) (*argW*) and tRNA^Arg^ (UCU) (*argU*) have unique anticodons, but *argW* is not essential because the rare AGG codon can also be decoded by wobble recognition by *argU* (49). The A35C mutation in *argVYZQ* and the T36G mutation in *argW* change the anticodon sequence from ACG or CCU to CCG, creating an additional tRNA^Arg^ (CCG) gene to supplement *argX*. Both *argVYZQ* and *argW* contain A37 rather than G37, so these recoded tRNAs would be unaffected by m^1^G37 deficiency. Interestingly, rapid adaptation of the tRNA pool through anticodon mutations has also been observed in response to tRNA gene deletion in yeast (50); our results show that perturbation of tRNA modifications is another potential driving force for recoding of tRNAs through anticodon mutations. In addition to anticodon mutations in *argVYZQ* and *argW*, two additional mutations, g(−8)a and g(−5)a, occur upstream of *argX* but downstream of the transcription start site at g(−13) (51), which might affect post-transcriptional processing of tRNA^Arg^ (CCG) by RNAse P (52). Importantly, the frequent occurrence of mutations in tRNA^Arg^ genes show that m^1^G37 deficiency affects translation of Arg codons as well as Pro codons.

Finally, three mutations in the C-terminal regulatory domain of *spoT* (P385R, S439R, and V593E) and a nonsense mutation in *ytfK* (L24*) are observed. These genes are associated with the stringent response, a stress response which is mediated by the alarmone (p)ppGpp and coordinates growth arrest under conditions of amino acid starvation, diverting cellular resources from protein synthesis and cell division to amino acid biosynthesis (40, 53). Mutations in *spoT* that disrupt its (p)ppGpp synthesis activity, decreasing intracellular (p)ppGpp concentrations, are commonly observed in experimental evolution studies as a generic mechanism to increase growth rate under conditions where the stringent response is activated (54). Similarly, the *spoT* mutations observed in response to m^1^G37 deficiency likely have the effect of decreasing the intracellular concentration of ppGpp to relax the stringent response, which might be activated in *trmD* mutant strains by uncharged tRNA^Pro^ or tRNA^Arg^, especially given that loss of m^1^G37 impairs tRNA^Pro^ aminoacylation.

## Discussion

Loss of the essential tRNA modification m^1^G37 is widely believed to cause cell death in bacteria due to an increase in the rate of translational frameshift errors at proline codons. In this work, we show in contrast that a single mutation in *proS* that restores aminoacylation of G37-unmodified tRNA^Pro^ is sufficient for substantial (>70%) recovery of the growth rate of m^1^G37-deficient *E. coli* strains. This result indicates that the fitness defects caused by deletion of *trmD* are mainly associated with impaired aminoacylation of tRNA^Pro^ rather than errors in protein translation. Although the pleiotropic effects of m^1^G37 deficiency (e.g., decreased expression of membrane proteins) have previously been attributed to the effects of translational frameshift errors at Pro codons, they can equally be attributed to the effects of diminished tRNA^Pro^ aminoacylation; indeed, consistent with this view, *trmD* knockdown strains of *E. coli* have a very similar phenotype to *proS* knockdown strains (13). Some similar conclusions based on an entirely different experimental approach have been reported in a recent preprint by Masuda et al. (55); in their work, ribosome profiling led to the discovery that loss of m^1^G37 induces A-site stalling of ribosomes at codons decoded by m^1^G37-containing tRNAs, which was shown to be mainly the result of defects in aminoacylation of unmodified tRNA^Pro^ and tRNA^Arg^ rather than ribosomal peptide bond formation. Consistent with this observation, overexpression of ProRS improved the growth of *trmD* knockdown strains. The effects of m^1^G37 deficiency on gene expression were also studied, showing activation of the stringent response in *trmD* knockdown strains. While the work of Masuda et al. provides a thorough systems-level analysis of the effects of m^1^G37 loss, our work provides detailed information on the evolutionary mechanisms available to counter m^1^G37 deficiency. Thus, the different strategies that were used to uncover the hidden functions of the m^1^G37 modification are highly complementary.

Although our results show that inefficient aminoacylation of tRNA^Pro^ is the main cause of growth defects observed in *trmD* knockdown strains, they do not imply that the m^1^G37 modification does not have a role in safeguarding the accuracy and efficiency of protein translation. Instead, it may be that this role of m^1^G37 is facilitated by the ability of ProRS to act as a gatekeeper of translational accuracy by discriminating between m^1^G37-modified and G37-unmodified tRNA^Pro^. If WT ProRS were unable to distinguish unmodified tRNA from mature tRNA, then unmodified tRNA could be aminoacylated and used as a substrate for protein synthesis, increasing the rate of frameshift errors in translation; this may explain why ProRS has evolved a preference for m^1^G37-modified tRNA. However, it should be noted that improvement of reading frame maintenance is a common feature of many tRNA modifications, including non-essential tRNA modifications (27); thus, it is possible that prevention of frameshift errors is a secondary function of m^1^G37. An alternative hypothesis is that m^1^G37 is used as a tRNA recognition element, ensuring selective aminoacylation of tRNA^Pro^ by ProRS and preventing aminoacylation by other aminoacyl tRNA synthetases. In any case, even though m^1^G37 is not indispensable for bacterial survival, the evolutionary persistence of *trmD* throughout the bacterial domain clearly indicates that it has some important role.

The rapid evolution of *E. coli* in response to m^1^G37 deficiency indicates that TrmD may be a more challenging antibiotic target than previously thought. *proS* has a large mutational target size, both because there are a large number of substitutions that increase aminoacylation efficiency of unmodified tRNA^Pro^ and because amplification of *proS* is another feasible adaptation to m^1^G37 deficiency, providing additional gene copies that can acquire mutations. Moreover, mutations in *proS* that increase the fitness of m^1^G37^−^ strains appear to have little impact on the growth of m^1^G37^+^ strains, indicating that similar polymorphisms could exist in natural populations of bacteria. These results indicate that bacteria challenged with an antibiotic targeting TrmD could readily evolve resistance through mutations in *proS* in addition to other general or target-specific mechanisms (e.g., increasing antibiotic efflux or mutations in *trmD* that disrupt antibiotic binding). Another potential problem is that there appears to be significant scope for increasing TrmD expression through a range of mutations upstream of *trmD*, due to the presence of strong mRNA secondary structure that inhibits translation; this might be another mechanism that can compensate for inhibition of TrmD. On the other hand, the observation that *trmD* mutant strains still show a fitness defect after ~200 generations of evolution and multiple rounds of adaptive mutations (Fig. 2) suggests that antibiotics targeting TrmD may show some benefit even if resistance does occur. However, direct antibiotic targeting of ProRS may be a potential strategy to induce the same advantageous phenotypic defects as found in *trmD* mutants (e.g., increased antibiotic susceptibility) without providing an additional pathway for the evolution of antibiotic resistance.

In this work, we showed that a recently developed approach to experimental evolutionary repair of essential genes (30) is a useful unbiased strategy to study the biological function of essential enzymes. In their original application of this method, Rodrigues and Shakhnovich showed that adaptation to loss of dihydrofolate reductase (DHFR) activity induced by catalytic mutations occurred mainly *via* loss-of-function mutations in other genes, which rerouted 2-deoxy-D-ribose-phosphate metabolism towards dTMP synthesis to compensate for the defect in DHFR. Thus, the evolutionary outcome was determined by the greater accessibility of loss-of-function mutations compared with rare compensatory mutations in the DHFR gene itself. We observed similarly that the evolutionary trajectory of *trmD* mutant strains was dictated by the high accessibility of mutations in *proS* that improve aminoacylation of unmodified tRNA. Whereas loss-of-function mutations frequently occur as an adaptive mechanism in evolutionary experiments (30, 35), our work provides a rare example where gain-of-function mutations are associated with a large mutational target, possibly due to the extensive protein-tRNA interaction surface of ProRS, which provides opportunity for a large number of mutations that could enhance the tRNA affinity of the enzyme (Fig. 5a). In contrast, in many cases, evolutionary adaptation of enzymes is highly constrained with few mutations available to improve enzyme activity (56, 57). Our work also emphasizes that the effect of gene amplification on mutational target size must also be considered to predict the outcome of evolutionary adaptation to inactivation of essential genes. In this case, amplification of *proS* provides another mechanism that increases the mutational target size of *proS* and thereby favors adaptation through mutations in *proS* rather than *trmD*. In contrast, amplification of *trmD* is not observed, most likely because the low translation efficiency of the *trmD* mRNA means that increasing the genomic copy number would not increase enzyme expression enough to outweigh the fitness cost associated with gene amplification. Thus, our results emphasize the importance of mutational target size in experimental evolution and show that this can be influenced by the balance between fitness costs and benefits associated with gene amplification.

Protein mistranslation has traditionally been viewed as a deleterious but inevitable phenomenon, minimized by the cell to the greatest extent practical (26). While mistranslation can indeed produce misfolded and dysfunctional proteins and have negative fitness effects, accumulating evidence suggests that organisms can tolerate errors in protein translation at surprisingly high frequencies, and in some cases, errors can be beneficial - a phenomenon known as adaptive translation (58). This view is consistent with our observation that evolutionary recovery of *E. coli* in response to m^1^G37 deficiency can occur without mutations that specifically address the increased rate of frameshift errors associated with unmodified tRNA^Pro^. While it is plausible that mutations in *proS* could have indirectly reduced the rate of frameshift errors by reducing stalling of ribosomes with Pro codons at the A-site, previous work has shown that m^1^G37 specifically prevents frameshift errors that occur during translocation of tRNA^Pro^ from the A-site to the P-site or during stalling of a Pro codon at the P-site (11, 27, 55). While frameshift errors are typically more deleterious than missense errors, they can also be harnessed as a regulatory mechanism, and can also unveil cryptic C-terminal sequences that may provide new protein functions. Indeed, TrmD is known to have at least one regulatory role in Mg^2+^ homeostasis, which is mediated by the effect of m^1^G37 on translation of Pro codons: a decrease in Mg^2+^-dependent TrmD activity activates transcription of the Mg^2+^ transporter *mgtA* in *Salmonella* by slowing translation of the Pro-rich leader sequence *mgtL* upstream of *mgtA* (59, 60). Our finding that the m^1^G37 modification is not indispensable for survival raises the possibility that TrmD may have additional dynamic roles in adaptive mistranslation beyond its housekeeping role in protein translation.

## Acknowledgements

B.E.C. was supported by a JSPS Postdoctoral Fellowship for Overseas Researchers from the Japan Society for the Promotion of Science. The project was funded by a KAKENHI Grant-in-Aid for Scientific Research (20F20705). Financial support from the Okinawa Institute of Science and Technology to P.L. is gratefully acknowledged. We thank Nobuhiko Tokuriki and Shimon Bershtein for critical reading of the manuscript, and Madhuri Gade for assistance with UPLC experiments.

## Author contributions

B.E.C. and P.L. designed the project, analyzed the data, and wrote the paper. B.E.C. and M.A.F. performed genome mutagenesis and purified the ProRS variants. G.U. isolated tRNA from *trmD* mutant strains for UPLC analysis. B.E.C. performed the remaining experimental work.

## Data availability

Sequence read data have been deposited to the NCBI Sequence Read Archive under BioProject PRJNA739405. Representative raw gel images from aminoacylation assays are provided in Supplementary File 1. The remaining data produced in this work are available upon request from the corresponding author.

## Materials and Methods

### General

*Escherichia coli* strain BW25113 was obtained from the National BioResource Project, National Institute of Genetics, Japan. The reference genome sequence of this strain was taken as GenBank sequence CP009273.1. A complete list of strains used in this study is given in Supplementary Table 6. Oligonucleotides were from Integrated DNA Technologies (Coralville, IA). Oligonucleotide sequences are listed in Supplementary Table 7. Synthetic DNA fragments were from Twist Bioscience (South San Francisco, CA) or Integrated DNA Technologies. Sanger sequencing was performed by FASMAC (Atsugi, Japan). A single batch of Luria-Bertani (LB) medium (Nacalai-Tesque, Kyoto, Japan) was used for experimental evolution and growth assays.

### Cloning

*TRM5* was cloned into the pBAD vector to facilitate protein expression under an arabinose-inducible promoter. The gene encoding TRM5 from *Saccharomyces cerevisiae* (UniProt: P38793), codon-optimized for expression in *E. coli* and cloned into the pET-28b(+) vector, was a gift from Dan Kozome. pBAD-EBFP2 was obtained from Addgene (plasmid #14891, courtesy of Robert Campbell). The *EBFP2* ORF in the pBAD-EBFP2 plasmid was replaced with the *TRM5* ORF in the pET-28b(+) vector (including the N-terminal hexahistidine and thrombin tags) by In-Fusion cloning using the In-Fusion HD Cloning Kit (Takara Bio, Kusatsu, Japan), yielding the pBAD-yTrm5 vector. For cloning of mutant *proS* variants into the pBAD vector, the *proS* gene was amplified either from wild-type *E. coli* BW25113 genomic DNA or population genomic DNA derived from the evolutionary experiment containing the desired mutation at a high frequency (>70%). The *TRM5* gene and thrombin tag in the pBAD-yTrm5 vector were replaced with the *proS* gene by In-Fusion cloning, yielding ProRS expression constructs with the N-terminal leader sequence MRGSHHHHHHSSG. To construct the tRNA^Pro^ (CGG) expression vector, the *proK* gene from *E. coli* BW25113, including the 8 bp flanking sequence on each side, was cloned into the EcoRI/PstI site of the pKK223-3 vector. This vector was assembled from the pBR322 vector (Promega, Madison, WI) and a synthetic DNA fragment by In-Fusion cloning. Successful assembly of each vector was confirmed by Sanger sequencing.

### Construction of mutant strains

Genomic mutations were introduced by multiplex automated genome engineering (MAGE) following literature protocols (61, 62). Mutagenic oligonucleotides were designed using the MODEST server (63). To construct the strains for experimental evolution, the five nucleotides immediately following the two consecutive stop codons of *glnH* were first mutated using a degenerate oligonucleotide to create a short barcode sequence (842706_842710 delins NNNNA). Growth assays were performed to confirm that the mutations introduced (including any potential off-target mutations) had no effect on growth rate (Supplementary Fig. 12). Mutations in *trmD* were then introduced into strains containing different barcode sequences.

Introduction of the barcode sequence was performed as follows. Electrocompetent BW25113 cells were co-transformed with the pORTMAGE-4 plasmid (Addgene plasmid #72679, courtesy of Csaba Pál) and the pBAD-yTrm5 plasmid. A single colony was transferred to 1.6 mL LB-Lennox medium containing chloramphenicol (25 μg/mL) and ampicillin (100 μg/mL) and incubated with shaking at 30 °C until the OD_600_ reached ~0.4. A 1 mL aliquot of the culture was transferred to a pre-heated 50 mL tube and incubated with shaking in a heat block at 700 rpm at 42 °C for 15 min. Cells were chilled on ice for 10 min, washed twice with ice-cold sterile water, and resuspended in 80 μL ice-cold sterile water. A 40 μL cell aliquot was transformed with 1 μL of 100 μM mutagenic oligonucleotide by electroporation (1.8 kV, 1 mm cuvette). The cells were transferred to 5 mL LB-Lennox medium and incubated with shaking at 30 °C for 1 h. Five milliliters of LB-Lennox containing 50 μg/mL chloramphenicol and 200 μg/mL ampicillin was added to the culture. After a further 4 h of incubation at 30 °C, 100 μL of a 10^4^ dilution of the culture was plated on an LB chlor amp (LB, 25 μg/mL chloramphenicol, 100 μg/mL ampicillin) agar plate and incubated overnight at 30 °C. Colonies containing the desired G to A mutation were identified using mutant allele-specific PCR (MASC-PCR) (61) and patched onto an LB chlor amp agar plate, which was incubated at 30 °C for several hours until growth was visible. The patches were struck onto fresh LB chlor amp agar plates and incubated at 30 °C overnight to obtain single colonies. Single colonies were used concurrently to create glycerol stocks, to repeat MASC-PCR to confirm the presence of the desired mutation, and to amplify a 750 bp fragment of genomic DNA surrounding the desired mutation, which was used to confirm the presence of a unique barcode sequence by Sanger sequencing. Genome mutagenesis of *trmD*, *proS*, *argV* and *argX* was achieved in the same way, except that in the case of *trmD* mutations, 0.1% (w/v) arabinose was added to the recovery medium 30 min after electroporation and included in all media thereafter to induce TRM5 expression.

### Plasmid curing

The pORTMAGE-4 plasmid was removed from the mutant strains by growth at 42 °C. Frozen cell stocks were struck onto LB amp ara (LB, 100 μg/mL ampicillin, 0.1% (w/v) arabinose) agar plates to obtain single colonies and incubated overnight at 30 °C. Single colonies were used to inoculate 1.6 mL LB amp ara, and the cultures were incubated for 1 h at 30 °C, then 4–5 h at 42 °C until turbid. One hundred microliters of a 10^4^ dilution of the culture was plated on an LB amp ara agar plate and incubated at 30 °C overnight. Single colonies were used to prepare glycerol stocks and were patched onto LB amp ara and LB amp ara + 25 μg/mL chloramphenicol agar plates to confirm chloramphenicol sensitivity.

The pBAD-yTrm5 plasmid was removed from the mutant strains using the CRISPR-Cas9 system pFREE (36). Electrocompetent cells were prepared from frozen cell stocks and transformed with the pFREE vector (Addgene plasmid #92050, courtesy of Morten Norholm). Transformed cells were plated on LB amp kan ara (LB, 100 μg/mL ampicillin, 50 μg/mL kanamycin, 0.1% (w/v) arabinose) agar plates and incubated overnight at 37 °C. A 1.6 mL aliquot of LB amp kan ara was inoculated with a single colony and incubated with shaking at 37 °C to mid-log phase (OD ~0.5). Five millilitres of LB containing rhamnose (0.2% (w/v)), anhydrotetracycline (200 ng/mL) and kanamycin (50 μg/mL) was inoculated with 5 μL of culture and incubated with shaking at 37 °C for 10 h to induce Cas9 and guide RNA expression. After induction, 100 μL of a 10^6^ dilution of the culture was plated on LB agar and incubated at 37 °C for 12–40 h until slow-growing colonies corresponding to cells that had lost the pBAD-yTrm5 plasmid (typically >95% of colonies) were visible. Colonies were purified by streaking the colonies onto a fresh LB agar plate and incubating at 37 °C for 12–48 h. Finally, a 1.6 mL aliquot of LB was inoculated with a single colony and incubated with shaking at 37 °C for 24 h. The resulting cultures were patched onto LB kan and LB amp ara plates to confirm antibiotic sensitivity and used to initialize growth assays and experimental evolution.

### Growth assays

To initialize growth assays from frozen cell stocks, cells were struck onto LB agar and incubated at 37 °C overnight. For each strain, at least two wells containing 100 μL LB medium in a 2 mL sterile deep-well 96-well plate were inoculated with a single colony. The plate was sealed with air-permeable film and incubated at 37 °C overnight with shaking at 1200 rpm. The next day, the overnight cultures were diluted into fresh LB medium in a 250 μL sterile round-bottom 96-well plate to give an OD_600_ of 0.002 ± 0.0005 in a volume of 150 μL. In order to minimize the effects of cross-contamination and evaporation, a checkerboard pattern was generally used, and edge wells were not used. The plate was covered with a lid treated with 0.05% (v/v) Triton X-100 in 20% ethanol to prevent condensation (64) and incubated at 37 °C with continuous shaking at 1200 rpm in a plate incubator. OD_600_ was measured periodically using a microplate reader. Each assay was performed at least in duplicate. Growth rates were estimated by linear regression of the linear portion of the baseline-subtracted curve of ln(OD_600_) versus time (approximately OD_600_ ~0.02 to 0.20).

### Experimental evolution

The Y86*, G117N, R154A, S165L and D169A *trmD* mutant strains were cured of the pBAD-yTrm5 plasmid as described above. Several replicate cultures, derived from different colonies obtained following induction of the pFREE vector, were prepared for each mutant strain (four cultures for Y86*, three cultures otherwise). These cultures were subjected to daily serial transfers for 15 days (99.7 generations) for the G117N and D169A mutants, 22 days (146.2 generations) for the R154A and S165L mutants, or 30 days (199.3 generations) for the Y86* mutant by transferring 200 μL culture to 20 mL fresh LB medium in a 100 mL flask and incubating the flask with shaking at 220 rpm at 37 °C for 24 h. Glycerol stocks and cell pellets for population genomic DNA extraction were frozen on days 1, 3, 6, 9, 15, 22, and 30. Growth assays were performed on days 0, 7, 15, 22 and 30. We initially planned to continue the experiment for 30 days; however, for the G117N and D169A mutants, the experiment was terminated after two consecutive growth assays showed wild-type growth rates for all replicate populations, while for the R154A and S165L mutants, the experiment was terminated upon temporary closure of the laboratory due to a typhoon.

### Next-generation sequencing

Population genomic DNA was extracted from frozen cell pellets using the Monarch Genomic DNA Purification Kit (New England Biolabs, Beverly, MA) according to the manufacturer’s instructions. Next-generation sequencing using the Illumina MiSeq platform (Illumina, San Diego, CA) was performed by the Okinawa Institute of Science and Technology DNA sequencing section (Onna, Okinawa, Japan) in two separate runs. In the first run, fifteen samples (3 × G117N populations at day 15, 3 × R154A populations at days 9 and 22, 3 × S165L populations at days 9 and 22) were sequenced in multiplex using the MiSeq v3 kit, generating 300 bp paired-end reads. In the second run, eight samples (4 × Y86* populations at days 9 and 30) were sequenced in multiplex using the MiSeq v2 kit, generating 150 bp paired-end reads. Adapter sequences were removed from the demultiplexed reads, and quality trimming was performed using bbduk version 38.87 in the BBTools package (65). Genomic mutations were identified by mapping the resulting reads onto the reference *E. coli* BW25113 genome sequence using breseq version 0.35.4 in polymorphism mode, using a frequency threshold of 5% to identify potential polymorphisms (66, 67). The sequencing results showed that one of the four Y86* populations was contaminated with *Micrococcus luteus*; this population was excluded from the analysis. Mean coverage depths ranged from 92.6 to 178.6 (mean 142.1) for each population in the first run and 126.0 to 134.3 (mean 129.0) for each population in the second run. The percentage of mapped reads was >98.6% for each population. Inversions, insertions, and duplications were identified through manual inspection of unassigned new junction evidence, and duplications were validated through analysis of read depth.

### Quantitative PCR

qPCR experiments were performed using a StepOnePlus Real-Time PCR System (Applied Biosystems, Bedford, MA). The mean copy number of *trmD* and *proS* in various population genomic DNA samples was measured using the relative standard curve method by comparison with the housekeeping genes *gyrB* and *tus*, which were assumed to remain at a single copy throughout the evolutionary experiment. Each well contained 2 ng of genomic DNA, 300 nM forward primer, 300 nM reverse primer, and 12.5 μL of 2x *Power* SYBR Green PCR Master Mix (Applied Biosystems) in a total volume of 25 μL. Thermocycling conditions were as follows: 95 °C for 10 min, then 30 cycles of 95 °C for 15 s and 60 °C for 60 s. Each plate contained negative controls and standards (wild-type *E. coli* BW25113 genomic DNA, 10-fold serial dilutions from 2 ng to 2 pg, in duplicate) to calculate efficiencies for each primer set. The mean amplification factors under the given reaction conditions were 1.91 (*trmD*), 1.95 (*proS*), 1.96 (*gyrB*), and 1.95 (*tus*). Primer specificity was validated by agarose gel electrophoresis and melt curve analysis. Data analysis was performed based on the method of Ganger et al. (68). The efficiency-weighted Δ*C_q_* values (Δ*C_q_*^(w)^) for each genomic DNA sample was calculated using the following equation: Δ*C_q_*^(w)^ = *C_q_*(gene of interest) × log_10_*E*(gene of interest) − 0.5 × (*C_q_*(*gyrB*) × log_10_*E*(*gyrB*) + *C_q_*(*tus*) × log_10_*E*(*tus*)), where *C_q_* is the threshold cycle value and *E* is the primer efficiency. One-way ANOVA with Dunnett’s test for multiple comparisons was used to assess the significance of differences in Δ*C_q_*^(w)^ values between the test samples and the wild-type control sample, using *P* < 0.05 as the significance threshold. The copy number of the gene of interest was then calculated as 10^(ΔΔ*C_q_*^(w)^), where ΔΔ*C_q_*^(w)^ is the difference in Δ*C_q_*^(w)^ values between the sample and the wild-type control.

### Isolation of mutant strains from evolved populations

To obtain double and triple mutant strains, glycerol stocks of *trmD* mutant populations from the evolutionary experiment were struck onto LB agar and incubated overnight at 37 °C to obtain single colonies. Populations were selected in order to minimize the number of loci that needed to be sequenced to confirm the genotype of the isolated strains, based on the NGS data (Supplementary Tables 3 and 6). The genomic region of interest was amplified by colony PCR and sequenced by Sanger sequencing. Strains containing the desired mutation were stored as glycerol stocks, and if necessary, sequencing of additional loci was performed to verify the genotype of the strain. For the majority of mutant strains, two strains derived from different colonies were isolated, in case additional mutations were acquired during processing and in case additional mutations were present in the population at a frequency too low to be detected by NGS.

### Identification of polymorphisms in *proS*

The *proS* gene was amplified from 50 ng population genomic DNA in a 25 μL PCR reaction using the high-fidelity polymerase PrimeSTAR Max DNA Polymerase (Takara Bio) and primers 134 and 138 (Supplementary Table 7). The thermocycling protocol was 30 cycles of 98 °C for 10 s, 55 °C for 5 s, then 72 °C for 10 s. The PCR product was purified using a NucleoSpin Gel and PCR Clean-up kit (Takara Bio), and Sanger sequencing was performed using primers 134, 135, 136, and 137 (Supplementary Table 7). Polymorphisms were identified by manual inspection of the sequencing chromatograms.

### Purification of bulk tRNA

Bulk tRNA was purified from various *E. coli* strains by phenol extraction, NaCl extraction, and isopropanol fractionation using a reported method (69). Cells were grown in 1 L 2×YT medium at 37 °C with shaking at 220 rpm to OD 1.2, chilled in an ice bath for 10 min, and centrifuged at 3000 × *g* for 12 min (4 °C). The cell pellets were washed once with 100 mL standard buffer (1 mM Tris-HCl pH 7.2, 10 mM MgCl_2_), flash frozen in liquid nitrogen, and stored at −80 °C until use. The cell pellets were thawed on ice and resuspended in 20 mL ice-cold standard buffer, and 30 mL of phenol saturated with standard buffer was added. The cell suspension was incubated on ice with frequent vortexing for 30 min, then centrifuged at 10000 × *g* for 15 min (15 °C). The upper aqueous phase was collected, and the organic phase was re-extracted with 20 mL standard buffer. Ethanol precipitation was performed by adding 0.1 volume of ice-cold 20% potassium acetate pH 5.0 and 2 volumes of ethanol (pre-chilled at −80 °C) to the combined aqueous extract and incubating at −80 °C overnight. The suspension was centrifuged at 10000 × *g* for 10 min (4 °C), the supernatant was discarded, and the tubes were inverted for several minutes to allow the ethanol to drain from the precipitate. The pellet was dissolved in 1 M NaCl and centrifuged at 12000 × *g* for 30 min (4 °C), and then the supernatant was collected and subjected to ethanol precipitation. The resulting pellet was dissolved in 10 mL of 0.5 M Tris-HCl pH 8.8, and the tRNA was deacylated by incubation at 37 °C with shaking for 1 h. Following another ethanol precipitation, the pellet was dissolved in 10 mL of ice-cold 0.3 M sodium acetate pH 7.0, and 0.54 volumes of ice-cold isopropanol was added dropwise with shaking between each addition. The mixture was centrifuged at 10000 × *g* for 7 min (4 °C), and the supernatant was set aside. The pellet was re-extracted by dissolving the pellet in 10 mL of 0.3 M sodium acetate pH 7.0 and repeating the isopropanol precipitation. The supernatants from each extraction were combined, and a further 0.44 volumes of isopropanol was added. Following centrifugation at 10000 × *g* for 10 min (4 °C), the pellet was washed with 10 mL of 70% ethanol (pre-chilled at −20 °C), drained of ethanol, and dissolved in a small volume of diethylpyrocarbonate-treated water (DEPC water). Aliquots were stored frozen at −80 °C. tRNA integrity and purity was confirmed by running a 5 μg aliquot on a 0.5% (v/v) bleach, 1.2% (w/v) agarose gel stained with ethidium bromide (70).

### Expression and purification of tRNA-Pro

Native m^1^G37 tRNA^Pro^ (CGG) was overexpressed from a pKK223-3 plasmid in *E. coli* BW25113 cells. Cells freshly transformed with the appropriate pKK223-3 plasmid by electroporation were grown in 1 L 2×YT medium at 37 °C with shaking at 220 rpm to OD ~0.5. tRNA expression was induced by addition of 0.5 mM isopropyl β-D-1-thiogalactopyranoside (IPTG), and the cultures were incubated for a further 2.5 h at 37 °C. Attempts to obtain the G37 tRNA^Pro^ (CGG) variant through overexpression of *proK* in *trmD* mutant strains were unsuccessful due to plasmid toxicity and low tRNA yields; thus, the G37 tRNA^Pro^ (CGG) variant was natively expressed in a fast-growing *trmD* mutant strain (BC_S159; *trmD* R154A, *proS* E19K, *argX* g(−5)a). In this case, cells were grown in 1 L 2×YT medium at 37 °C with shaking at 220 rpm to OD ~1.2. In both cases, the cells were harvested, and bulk tRNA extraction was performed as described above. tRNA^Pro^ (CGG) was purified by hybridization to a DNA oligonucleotide immobilized on a streptavidin resin following a literature protocol (71). Briefly, a biotinylated DNA oligonucleotide complementary to the target tRNA (Supplementary Table 7) was immobilized on Novagen Streptavidin Agarose resin (Merck-Millipore, Burlington, MA) in a 0.22 μm Ultrafree-MC filter cup (Merck-Millipore) (1.5 nmol oligonucleotide per 100 μL resin). The resin was equilibrated with 10 mM Tris (pH 7.6) and incubated with bulk tRNA (800 μg per 100 μL resin) dissolved in hybridization buffer (10 mM Tris, 0.9 M NaCl, 0.1 mM EDTA, pH 8.0) at 65 °C for 10 min. The resin was washed with 10 mM Tris pH 7.6 until the A260 of the flow-through was < 0.01. The target tRNA was eluted twice by incubation of the resin in 10 mM Tris (pH 7.6) at 65 °C for 5 min, followed by rapid centrifugation (10000 × *g*, 30 s). The tRNA was ethanol precipitated, washed with 70% ethanol, dissolved in 10 mM Tris, 10 mM MgCl_2_ (pH 7.6), and stored in aliquots at −80 °C. tRNA integrity and purity was confirmed by running a 20 ng aliquot on a 7 M urea / 10% polyacrylamide gel stained with SYBR Gold.

### Expression and purification of ProRS

Wild-type *E. coli* ProRS was overexpressed in *E. coli* BW25113 cells. The E19K, G200S, and E514V variants of *E. coli* ProRS were overexpressed in BW25113 variants that contained the same mutation in the chromosomal copy of *proS* in order to exclude the possibility of contamination with the wild-type enzyme (strains BC_S171, BC_S173, and BC_S176, respectively; Supplementary Table 6). Cells freshly transformed with the relevant expression vector by electroporation were grown at 37 °C with shaking at 220 rpm in 1 L Terrific Broth (TB) medium to OD_600_ ~0.6, and ProRS expression was induced with 0.2% (w/v) arabinose. The cultures were incubated with shaking at 30 °C for a further 20 h. The cells were harvested by centrifugation, resuspended in lysis buffer (20 mM Tris, 0.5 M NaCl, 20 mM imidazole, 5 mM MgCl_2_, 2 mM β-mercaptoethanol, pH 8.0), lysed by sonication, and fractionated by centrifugation at 10000 × *g* for 1 h (4 °C). The soluble fraction of the cell lysate was sonicated again to fragment genomic DNA, then filtered with a 0.45 μm syringe filter and loaded onto a 1 mL HisTrap HP 1 mL column (Cytiva, Marlborough, MA) equilibrated with lysis buffer. The column was washed with 5 volumes of lysis buffer, 10 volumes of wash buffer 1 (20 mM Tris, 1 M NaCl, 5 mM MgCl_2_, 2 mM β-mercaptoethanol, pH 8.0), and 10 volumes of wash buffer 2 (20 mM Tris, 0.5 M NaCl, 32 mM imidazole, 5 mM MgCl_2_, 2 mM β-mercaptoethanol, pH 8.0). The protein was then eluted from the column with 5 volumes of elution buffer (20 mM Tris, 0.5 M NaCl, 0.5 M imidazole, 5 mM MgCl_2_, 2 mM β-mercaptoethanol, pH 8.0), concentrated to 400 μL using a 10 KDa molecular weight cut-off centrifugal filter, and loaded onto a Superdex 200 Increase 10/300 column (Cytiva) equilibrated with storage buffer (20 mM HEPES, 0.15 M NaCl, 5 mM MgCl_2_, 5 mM β-mercaptoethanol, pH 7.50). The protein was eluted from the size-exclusion column in storage buffer at a rate of 0.75 mL/min and the peak corresponding to dimeric ProRS was collected. Glycerol was added to the protein to a final concentration of 10%, and the protein concentration was adjusted to 1 mg/mL. The protein was flash-frozen in small aliquots (<100 μL) in liquid nitrogen and stored at −80 °C. Protein purity was confirmed by SDS-PAGE.

### Aminoacylation assays

Aminoacylation assays were performed with saturating ATP and L-proline using a gel shift assay based on selective biotinylation of aminoacylated tRNA using an NHS-biotin reagent, followed by streptavidin conjugation and separation of conjugated and unconjugated tRNA by urea PAGE (72). For comparison of enzyme activity between ProRS variants, the assay was performed as follows. The m^1^G37 and G37 variants of tRNA^Pro^ (CGG) were diluted to 8 μM in refolding buffer (10 mM Tris, 10 mM MgCl_2_, pH 7.6 in RNAse-free water) and refolded by heating at 85 °C for 2 min, then 37 °C for 15 min. ProRS variants were diluted to 6 μM and 15 nM in 1× assay buffer (50 mM Tris pH 7.6, 20 mM KCl, 4 mM DL-dithiothreitol, 10 mM MgCl_2_, 4 mM ATP, 2.5 mM L-proline, 0.1 mg/mL bovine serum albumin in RNAse-free water). Reaction mixtures were prepared by mixing 9 μL 2× assay buffer, 1.5 μL RNAse-free water and 4.5 μL tRNA and pre-heated at 37 °C for 3 min. The reaction was initiated by the addition of 3 μL of 15 nM enzyme, giving a final reaction volume of 18 μL, tRNA concentration of 2 μM, and enzyme concentration of 2.5 nM, and the reaction mixture was incubated at 37 °C. At the 7 min and 14 min time points, 5 μL aliquots of the reaction mixture were quenched by addition of 7.5 μL quench buffer (0.42 M sodium acetate (pH 5.0), 16.7 mM EDTA, 0.13 mg/mL glycogen) followed by 37.5 μL ethanol, and stored at −20 °C. The control reactions were processed similarly, except that the final tRNA concentration was 0.5 μM, the final enzyme concentration was 1 μM for the positive controls, and a 10 μL aliquot of the reaction mixture was quenched after 14 min. After completion of the assay, the tubes were transferred to −80 °C and incubated for 30 min. The precipitated tRNA was collected by centrifugation at 21,000 × *g* (10 min, 4 °C), washed twice with cold 70% ethanol, and dissolved in DEPC water (16 μL for test samples, 8 μL for controls). A ~40 mM solution of EZ-Link sulfo-NHS-biotin in 60 mM HEPES (pH 8.0) was prepared. Four microliters of this solution was mixed with 4 μL of tRNA solution, centrifuged briefly, and incubated on ice for 2 h. The tRNA was ethanol precipitated by addition of 0.8 μL 20% potassium acetate (pH 5.0) and 24 μL ethanol followed by incubation at −80 °C for 30 min. The precipitate was collected by centrifugation at 21,000 × *g* (10 min, 4 °C), washed once with cold 70% ethanol, and dissolved in 10 μL DEPC water. The ethanol precipitation and 70% ethanol wash was repeated, and the tRNA was then dissolved in 8 μL DEPC water. Four microliters of this solution was mixed with 1 μL of 1 mg/mL streptavidin and incubated at room temperature for 20 min. Five microlitres of 2× RNA loading dye (New England Biolabs) was added, and the sample was loaded without heating onto a 7 M urea 10% polyacrylamide gel and run at 80 V for 10 min, followed by 200 V for 50 min. The gel was stained with SYBR Gold (Sigma, St Louis, MO) for 25 min, then imaged using an iBright FL1500 imaging system (Thermo Fisher Scientific). Because of the large excess of streptavidin used, streptavidin-conjugated aminoacylated tRNA appeared as a single band. The percentage of aminoacylated tRNA in each sample was calculated by band integration in ImageJ version 1.53a. Experiments to determine Michaelis-Menten kinetic parameters were performed similarly, except that the tRNA concentration was varied, and the enzyme concentration was adjusted to ensure linearity of product formation over the course of the assay. The volume of DEPC water used to redissolve the tRNA pellet before biotinylation was adjusted to give a maximum of 2.5 pmol of tRNA in the biotinylation reaction. For data analysis, the product concentration in each sample was taken as the fraction of aminoacylated tRNA determined by band integration multiplied by the total tRNA concentration. The substrate concentration was taken as the concentration of aminoacylation-competent tRNA, which was determined by multiplying the total tRNA concentration by the fraction of aminoacylated tRNA in the positive control. For determination of Michaelis-Menten parameters, the initial rate of product formation was plotted against the substrate concentration and fitted to the substrate inhibition model (for m^1^G37-modified tRNA) or a linear model (for G37-unmodified tRNA, to estimate *k*_cat_/*K*_M_) in GraphPad Prism version 8.3.0.

### Analysis of tRNA m1G content

tRNA m^1^G content was determined by ultra-performance liquid chromatography (UPLC) analysis of a nucleoside digest. Purified tRNA^Pro^ (CGG) (2 μg) was heated at 85 °C for 2 min, chilled immediately on ice, and then digested with 2 μL Nucleoside Digestion Mix (#M0649, New England Biolabs) in a 50 μL reaction incubated at 37 °C overnight. After digestion, 50 μL ethanol was added and the mixture was centrifuged at 21,000 × *g* for 10 min. The supernatant was transferred to a clean microcentrifuge tube and evaporated to dryness in a vacuum evaporator. The residue was dissolved in 50 μL of 2 mM ammonium formate, filtered through a 0.22 μm filter, and analyzed by UPLC. The separation was performed using a ACQUITY UPLC BEH C18 column (1.7 μm pore size, 2.1 × 100 mM) on a Waters ACQUITY H-Class UPLC system (Waters, Milford, MA) equipped with a photodiode array detector (ACQUITY UPLC PDA eλ detector, Waters) and mass detector (ACQUITY UPLC qDa detector, Waters). The following parameters were used: injection volume 3 μL, flow rate 0.2 mL/min, column temperature 40 °C, buffer A = 2 mM ammonium formate, buffer B = 90% acetonitrile, 10% water, 0.1% formic acid. The following program was used for elution: hold at 2% B to 2 min, gradient to 5% B to 8 min, hold at 5% B to 13 min, gradient to 10% B to 19 min, hold at 10% B to 24 min, gradient to 30% B to 30 min, hold at 60% B to 31 min, then hold at 2% B to 47 min. Parameters for mass detection were as follows: spectral range 100-600 Da, positive mode electrospray ionization, cone voltage 15 V, capillary voltage 0.8 kV. The following nucleosides were used as analytical standards: adenosine (>98%, 011-24593; FUJIFILM Wako Pure Chemical Co., Osaka, Japan), cytosine (>98%, 031-23233; FUJIFILM Wako Pure Chemical Co.), guanosine (>98%, G6752; Sigma), pseudouridine (>98%, NP11297; Carbosynth, Compton, United Kingdom), uridine (>98%, U0020; Tokyo Chemical Industry Co., Tokyo, Japan), *N*^1^-methylguanosine (>98%, NM08574; Carbosynth).

