## Supplementary Information for "Evolutionary repair reveals an unexpected role of the tRNA modification m^1^G37 in aminoacylation"

**Supplementary Figure 1. Growth rates of key *E. coli* strains and evolved populations.** For each strain or population, the mean and individual replicate values are shown, except in panel **h** where mean ± s.d. is shown (*n* = 12 to 24). The dotted lines indicate the growth rate of the WT strain (1.53 $h^{-1}$) and the mean growth rate of the R154A, S165L, D169A, and Y86* strains (0.54 $h^{-1}$). The precise number of replicates for each strain is indicated in Figures 1-3. **a,** Strains with mutations in *trmD* (Fig. 1b). **b-f**, Evolving populations (Fig. 2). **b**, G117N; **c**, S165L; **d**, D169A; **e**, R154A; **f**, Y86*. **g**, Mutants of the *trmD* G117N strain with a second mutation in *trmD* (Fig. 3b). **h**, Mutants of the *trmD* R154A strain with a single mutation in *proS* (Fig. 3c and Supplementary Fig. 5). **i**, Mutants of the *trmD* R154A strain with mutations in *argV* and *argX* (Fig. 3d).

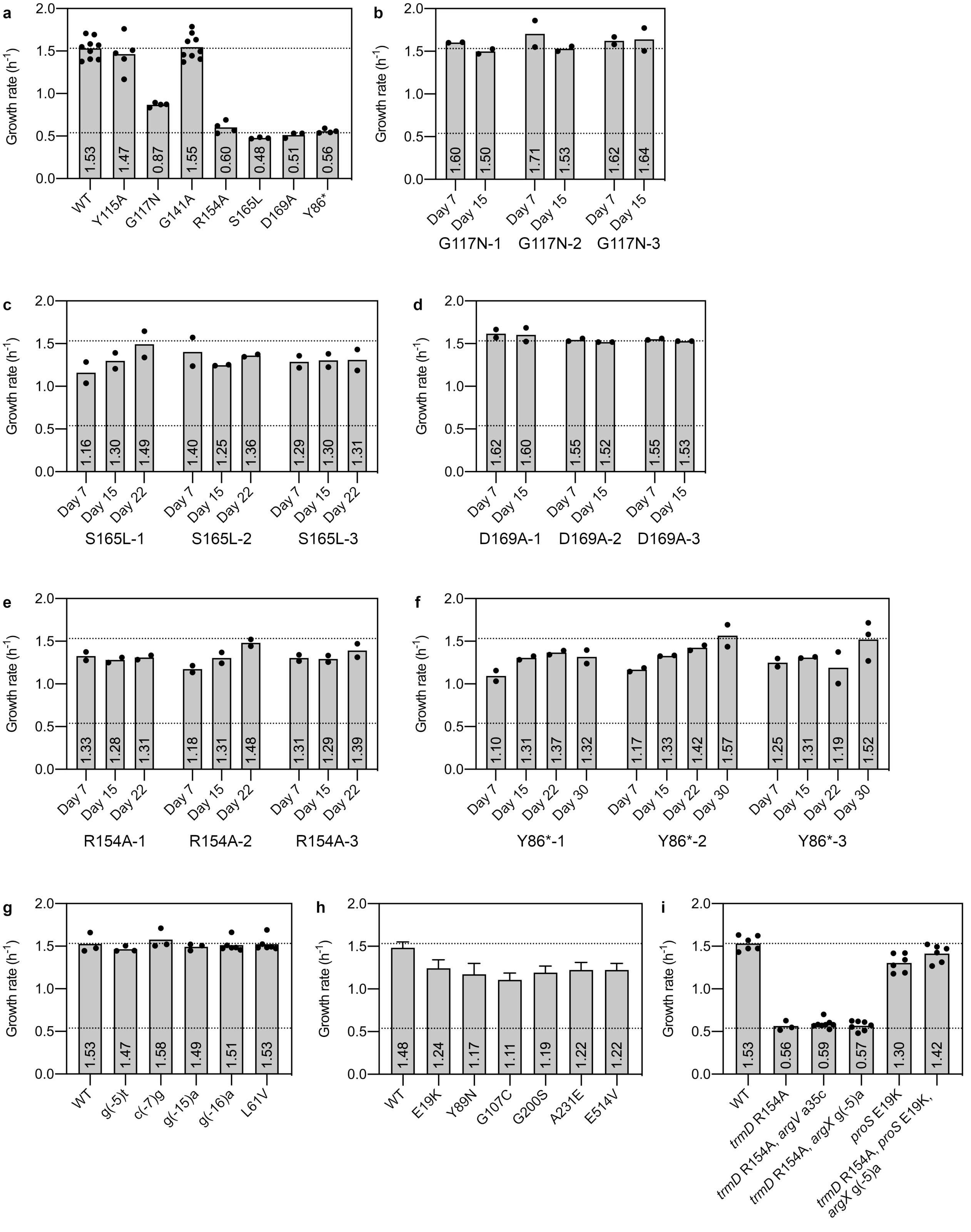
a
Growth rate (h-1)
2.0
1.5
1.0
0.5
0.0
1.53
1.47
0.87
1.55
0.60
0.48
0.51
0.56
WT
Y115A
G117N
G141A
R154A
S165L
D169A
Y86*
b
1.60
1.50
1.71
1.53
1.62
1.64
Day 7
Day 15
G117N-1
G117N-2
G117N-3
c
1.16
1.30
1.49
1.40
1.25
1.36
1.29
1.30
1.31
Day 22
S165L-1
S165L-2
S165L-3
d
1.62
1.60
1.55
1.52
1.55
1.53
D169A-1
D169A-2
D169A-3
e
1.33
1.28
1.31
1.18
1.31
1.48
1.31
1.29
1.39
R154A-1
R154A-2
R154A-3
f
1.10
1.31
1.37
1.32
1.17
1.33
1.42
1.57
1.25
1.31
1.19
1.52
Day 30
Y86*-1
Y86*-2
Y86*-3
g
1.53
1.47
1.58
1.49
1.51
1.53
WT
g(-5)t
c(-7)g
g(-15)a
g(-16)a
L61V
h
1.48
1.24
1.17
1.11
1.19
1.22
1.22
WT
E19K
Y89N
G107C
G200S
A231E
E514V
i
1.53
0.56
0.59
0.57
1.30
1.42
WT
trmD R154A
trmD R154A, argV a35c
trmD R154A, argX g(-5)a
proS E19K
trmD R154A, proS E19K, argX g(-5)a

**Supplementary Figure 2. UPLC analysis of nucleoside content in *trmD* mutant strains. a,** Separation of nucleoside standards. Left panel, UV chromatogram with detection at 260 nm; right panel, extracted ion chromatogram for $m^1G$ ($[M+H]^+$ $m/z$ 298). Abbreviations: A, adenosine; C, cytosine; G, guanosine; U, uridine; $m^1G$, $N^1$-methylguanosine; ψ, pseudouridine. **b**, Separation of digested $tRNA^{Pro}$ (CGG) purified from various *trmD* mutant strains of *E. coli*. Left panel, UV chromatogram with detection at 260 nm; right panel, extracted ion chromatogram for $m^1G$ ($[M+H]^+$ $m/z$ 298). The elution time of $m^1G$ (16.198 min) is indicated by a dashed line. Data for the WT and Y86* strains are reproduced from Fig. 1d for comparison with the other strains. **c**-**e**, Evaluation of assay sensitivity. Digested WT $tRNA^{Pro}$ (CGG) was spiked into a mixture of 20 μM each of adenosine, cytosine, guanosine and uridine at the indicated concentration. **c,** $A_{260}$ chromatogram of a 5% (v/v) dilution of digested tRNA. **d,** Extracted ion chromatogram of a 1% (v/v) dilution of digested tRNA. **e,** Extracted ion chromatogram for digested $tRNA^{Pro}$ (CGG) from the *trmD* Y86* strain, showing that $m^1G$ is not detected with a limit of detection <1%.

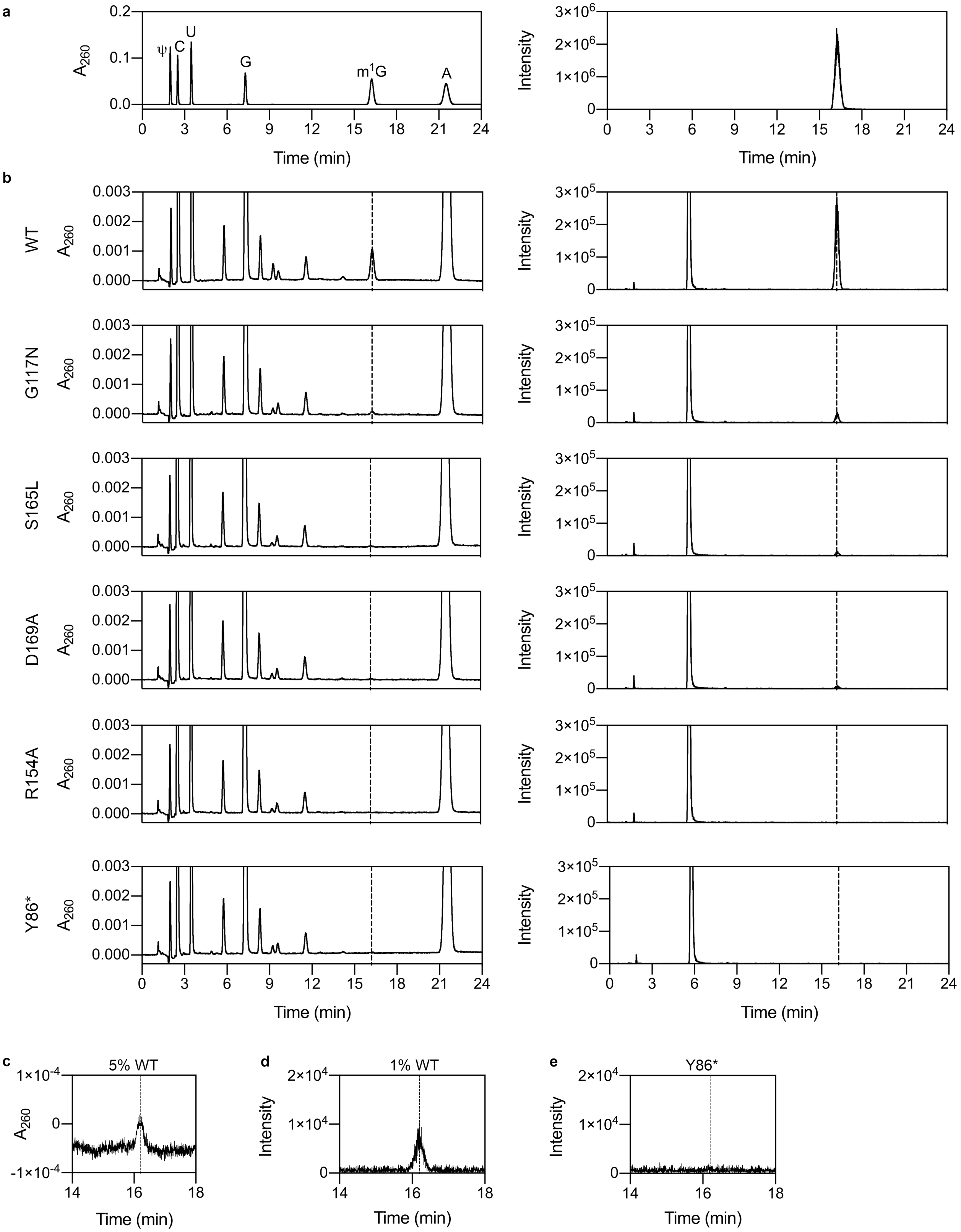
a
b
c
d
e
A260
Intensity
Time (min)
ψ
C
U
G
m1G
A
WT
G117N
S165L
D169A
R154A
Y86*
5% WT
1% WT

**Supplementary Figure 3. Copy number of *trmD* in evolving populations.** Mean copy number of *trmD* in population genomic DNA extracted from the G117N, S165L, D169A and R154A populations at different time points was measured by qPCR. Error bars represent 95% confidence intervals. Estimates are derived from two technical replicates per primer pair for each sample.

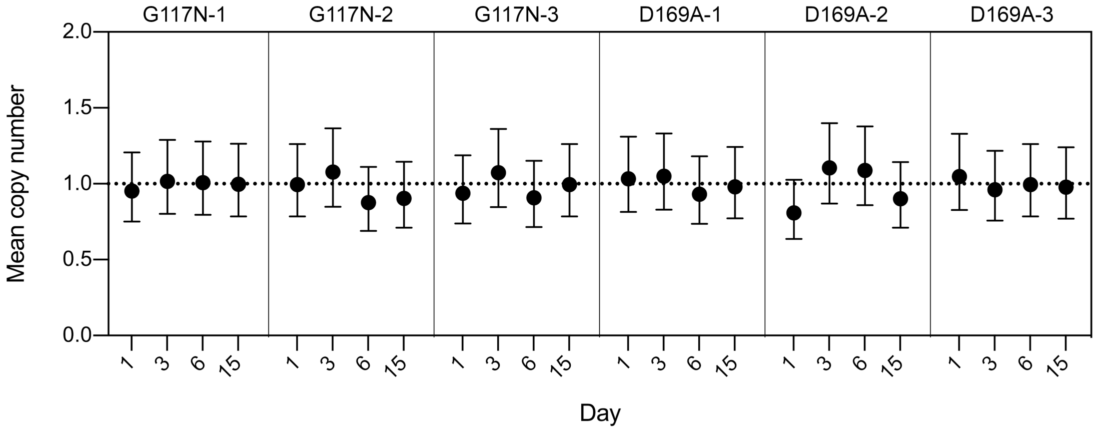


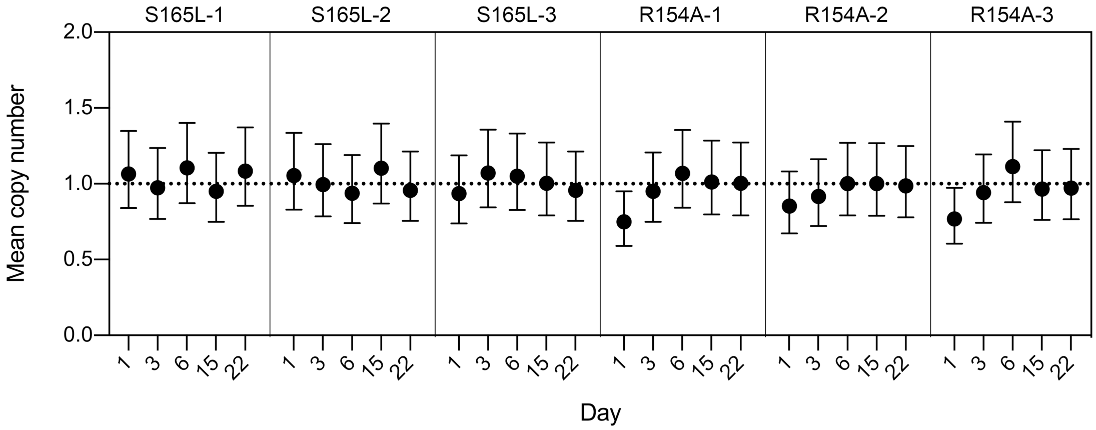

**Supplementary Figure 4. Off-target mutations introduced during MAGE do not affect growth of *trmD* mutant strains.** Growth assays of each strain transformed with the pBAD-yTrm5 and pFREE plasmids (i.e., the final strains to be stored as a glycerol stock before the evolutionary experiment) were performed in LB media containing 25 µg/mL kanamycin, 100 µg/mL ampicillin and 0.2% (w/v) glucose at 37 °C. Data represent mean ± s.d. of two biological replicates (different colonies).

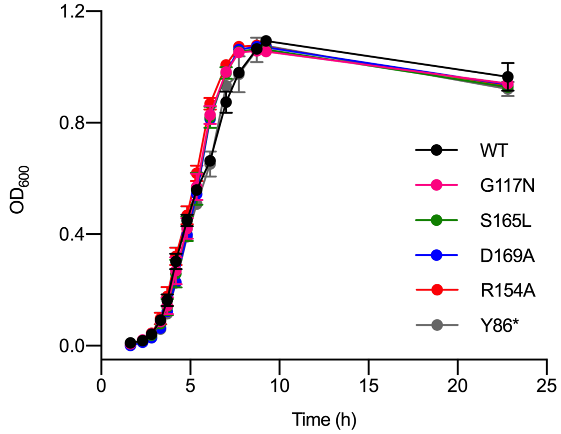

**Supplementary Figure 5. Replicate growth assays of variants of the *trmD* R154A strain with an additional mutation in *proS*.** Data represent mean ± s.d. WT, E514V, *n* = 3 colonies; G200S, A231E, E19K, Y89N, G107C, *n* = 6 colonies (three colonies each from two independently isolated strains). The E19K, A231E, and E514V variants show consistently faster growth than the Y89N, G107C, and G200S variants when the full growth curve is considered.

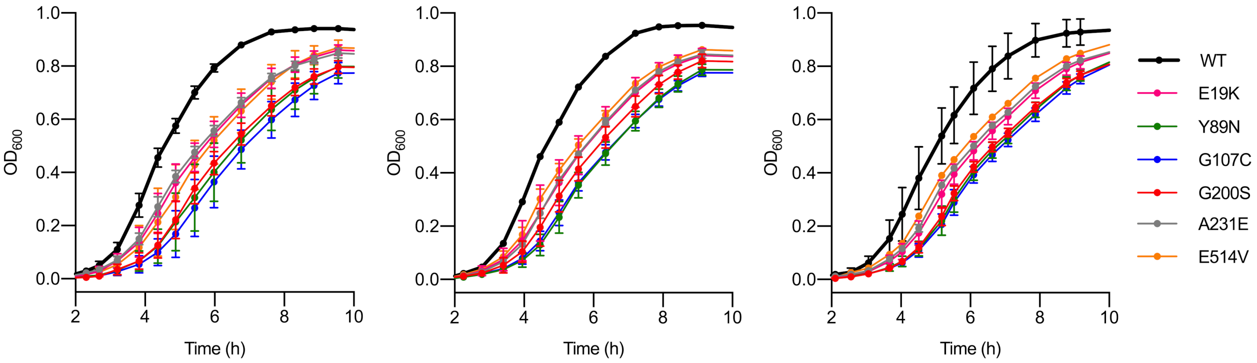

**Supplementary Figure 6. Sanger sequencing analysis of *proS* at early time points in the evolutionary experiment.** The *proS* gene was amplified from population genomic DNA at each time point for the S165L, R154A, and Y86* populations, and polymorphisms were identified by Sanger sequencing. Each sequencing trace shows three bases centered on the mutated base (not necessarily the mutated codon). Mutations are described relative to the antisense strand. The frequency of each mutation at day 9, determined by NGS, is also indicated.

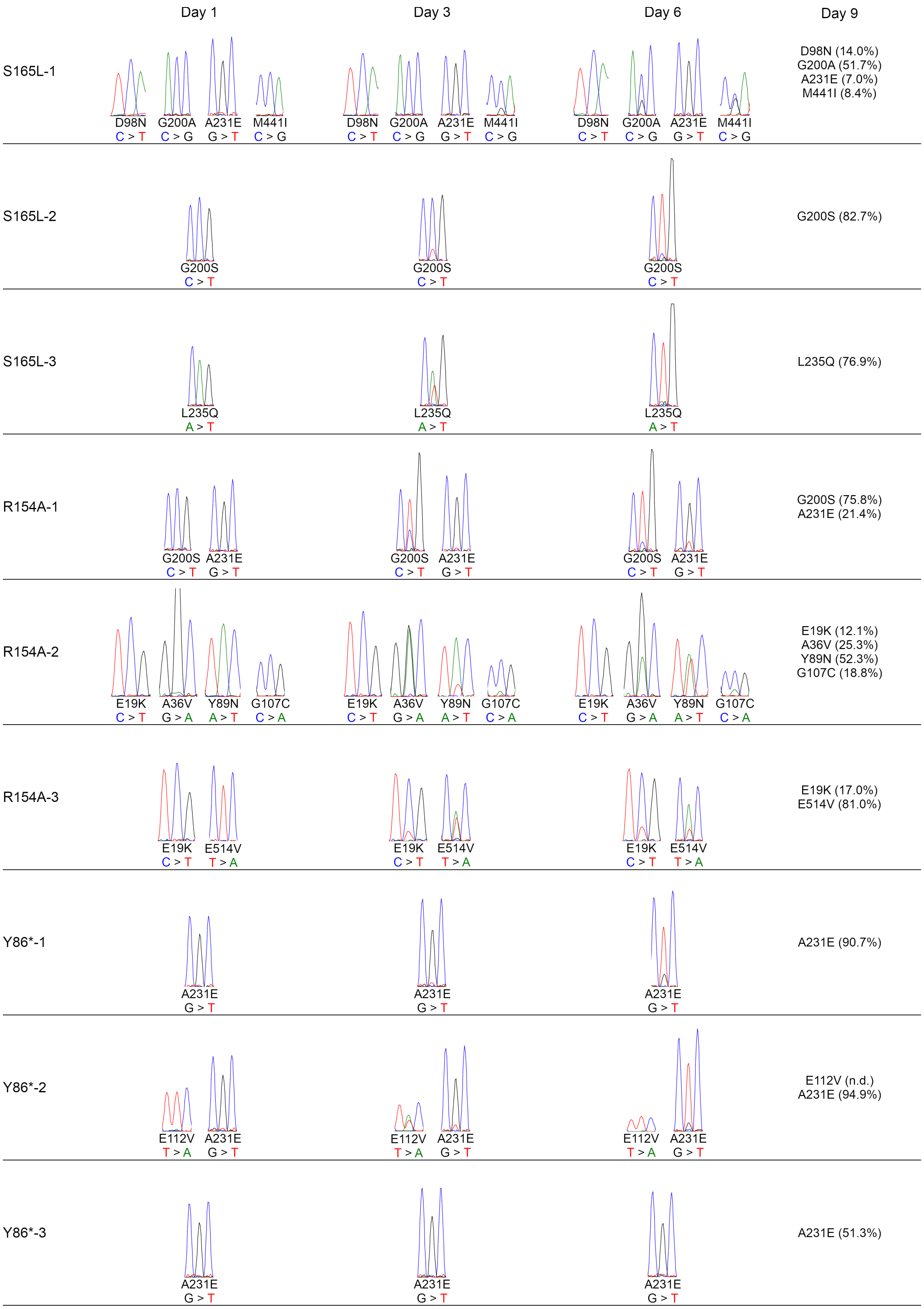
Day 1
Day 3
Day 6
Day 9
S165L-1
D98N C > T
G200A C > G
A231E G > T
M441I C > G
D98N (14.0%)
G200A (51.7%)
A231E (7.0%)
M441I (8.4%)
S165L-2
G200S C > T
G200S (82.7%)
S165L-3
L235Q A > T
L235Q (76.9%)
R154A-1
G200S C > T
A231E G > T
G200S (75.8%)
A231E (21.4%)
R154A-2
E19K C > T
A36V G > A
Y89N A > T
G107C C > A
E19K (12.1%)
A36V (25.3%)
Y89N (52.3%)
G107C (18.8%)
R154A-3
E19K C > T
E514V T > A
E19K (17.0%)
E514V (81.0%)
Y86*-1
A231E G > T
A231E (90.7%)
Y86*-2
E112V T > A
A231E G > T
E112V (n.d.)
A231E (94.9%)
Y86*-3
A231E G > T
A231E (51.3%)

**Supplementary Figure 7. Duplication of *proS* observed in the Y86*-3 population at day 9. a,** Boundaries of the duplicated region identified by new junction analysis. **b,** Validation of duplicated region by analysis of read coverage depth. The light gray region corresponds to the duplicated region, and the dark gray region corresponds to the *proS* open reading frame. The dotted line represents the mean read coverage depth across the whole genome.

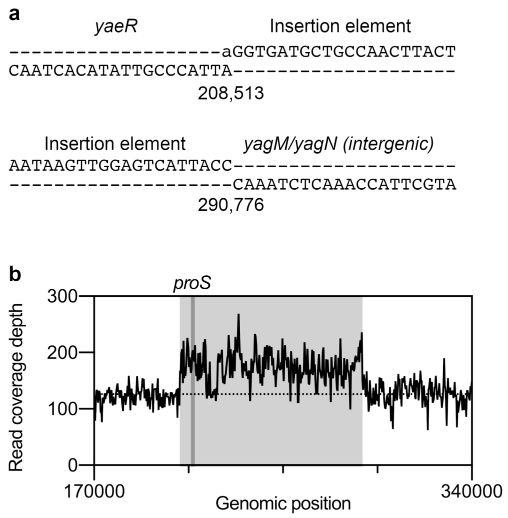

**Supplementary Figure 8. Gel shift analysis of the aminoacylation activity of ProRS variants on m$^1$G37-modified and G37-unmodified tRNA$^{Pro}$ (CGG).** Reaction conditions: tRNA concentration 2 µM, enzyme concentration 2.5 nM, reaction temperature 37 °C, reaction time 14 min. The enzyme concentration was chosen to ensure linearity of product formation for the WT enzyme with m$^1$G37-modified tRNA over the course of the assay. + (1), positive control with 1 µM WT ProRS; + (2), positive control with 1 µM ProRS E514V; -, no enzyme control. The percentage of aminoacylated tRNA in each lane is indicated. Each panel (**a**-**c**) shows a single experimental replicate. The replicate shown in Fig. 5b corresponds to panel **c**. Raw gel images are provided in Supplementary Data 1.

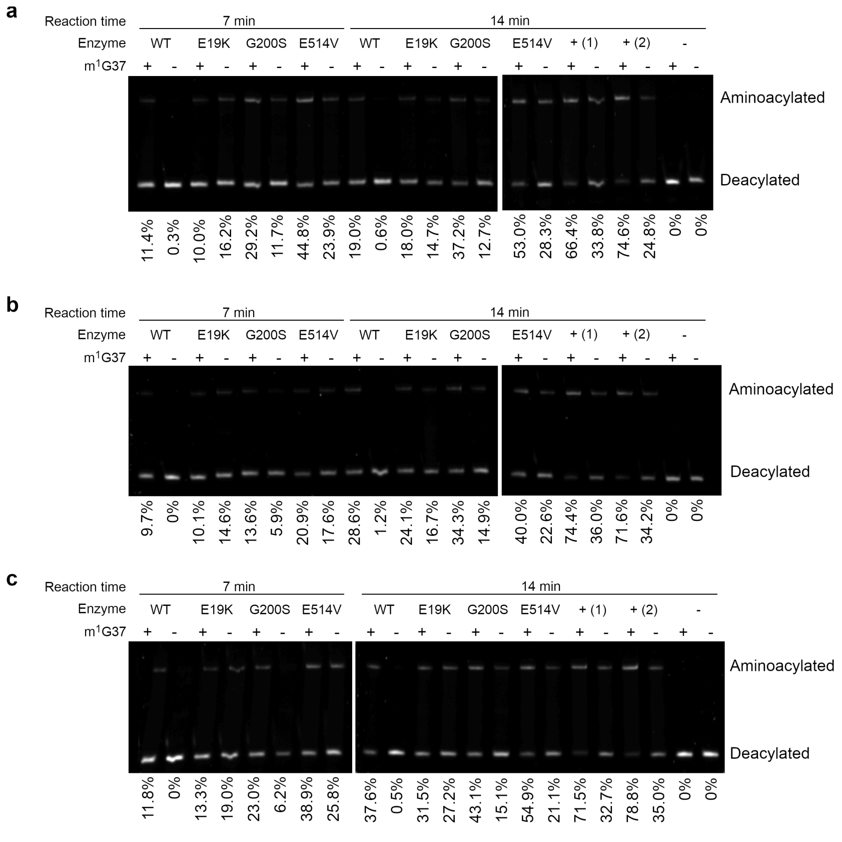

**Supplementary Figure 9. Gel shift analysis for determination of Michaelis-Menten parameters for aminoacylation of m$^1$G37-modified and G37-unmodified tRNA$^{Pro}$ (CGG) by WT and E514V ProRS.** The rate of aminoacylation was measured using a 2-fold dilution series of tRNA. One representative experimental replicate (each with two technical replicates in the case of m$^1$G37-modified tRNA) is shown for each enzyme-tRNA combination. The percentage of aminoacylated tRNA in each lane is indicated. Raw gel images are provided in Supplementary Data 1. Abbreviations: n.q., not quantified; (+), positive control with 1 µM enzyme; (-), negative control with no enzyme. **a,** WT ProRS + m$^1$G37 tRNA. Enzyme concentration, 0.8 nM; maximum total tRNA concentration 6.29 µM. **b,** E514V ProRS + m$^1$G37 tRNA. Enzyme concentration, 0.267 nM; maximum total tRNA concentration 4.15 µM. **c,** WT and E514V ProRS + G37 tRNA. Enzyme concentration, 15 nM (WT) or 1 nM (E514V); maximum total tRNA concentration 3.70 µM.

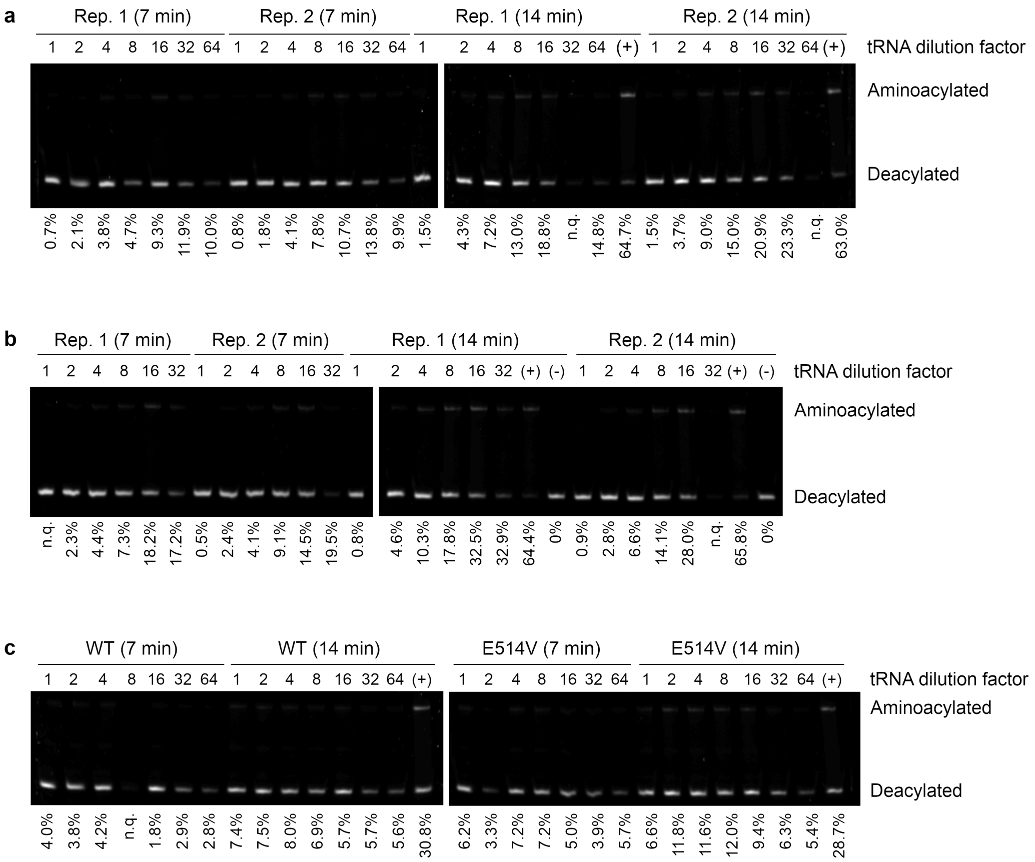

**Supplementary Figure 10. Mutations in *proS* do not affect growth of *E. coli* BW25113.** Growth assays of the WT BW25113 strain and variants with mutations in *proS* (E19K, G200S, or E514V) in LB media at 37 °C. Data represent mean ± s.d.; WT, *n* = 3 colonies; *proS* mutants, *n* = 6 colonies (three colonies each from two independently isolated strains).

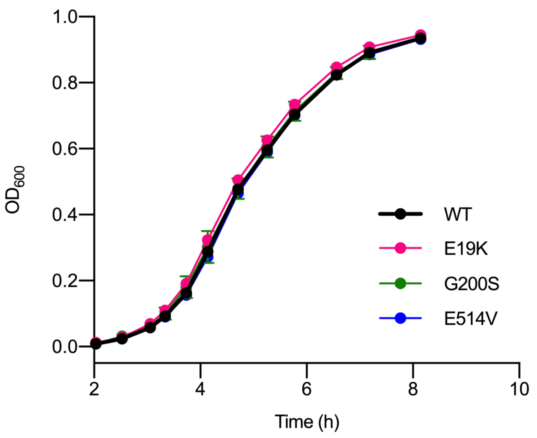

**Supplementary Figure 11. Effect of mutations upstream of the *trmD* ORF on predicted mRNA secondary structure. a,** Structure of *trmD* mRNA from the -30 position (immediately following *rimM*) to the +28 position, predicted using RNAfold version 2.4.18 (Lorenz et al., 2011). Positions where a mutation is observed in the evolutionary experiment are shown in light blue, and the start codon of *trmD* is shown in dark blue. The mRNA structure was visualized using Forna (Kerpedjiev et al., 2015). **b,** Effect of point mutations on predicted minimum free energy of *trmD* mRNA secondary structure (positions -30 to +28). Free energy calculations were performed using RNAfold version 2.4.18 (Lorenz et al., 2011).

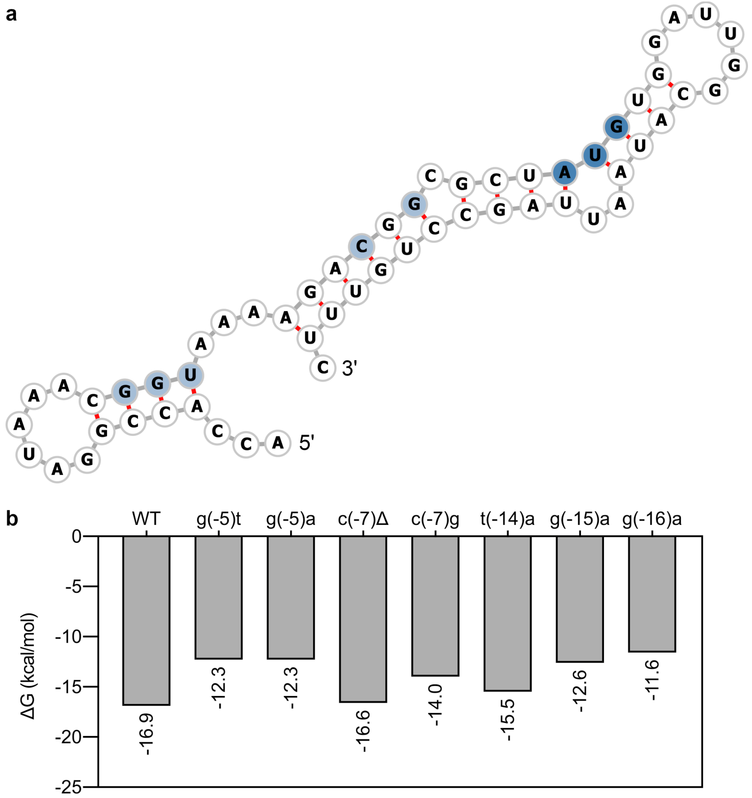

**Supplementary Figure 12. Barcode mutations do not affect growth of *E. coli* BW25113.** Growth assays of the WT BW25113 strain (black) and the seven barcoded strains (gray; see Supplementary Table 6) in LB media at 37 °C. Data represent mean ± s.d. of three biological replicates (different colonies).

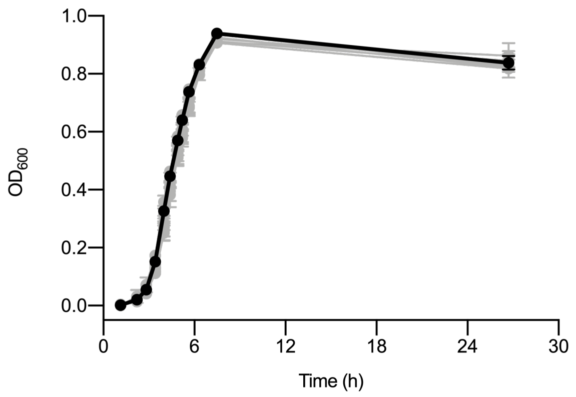

**Supplementary Table 1. Previously reported effects of TrmD substitutions on catalytic activity and bacterial growth.**

| Substitution | Effect on catalytic activity | Effect on growth | Notes |
|---|---|---|---|
| Y115A | 8-fold decrease in $k_{cat}/K_M$ for tRNA and 12-fold decrease in $k_{cat}/K_M$ for SAM (Elkins et al., 2003).<br><br>6-fold increase in $K_d$ for SAM, 34-fold increase in $K_d$ for tRNA, and 22-fold decrease in $k_{chem}$ (Christian et al., 2016). | Unknown. | - |
| G117N | Unknown; however, G117A abolishes activity (Elkins et al., 2003). | Reduces colony size to <10% of wild-type strain in *Salmonella enterica* Typhimurium (37 °C) (Björk et al., 2001). | G117S and G117Q *trmD* mutants of *S. enterica* Typhimurium are viable (Björk et al., 2001). |
| G141A | 1400-fold decrease in $k_{cat}/K_M$ for SAM (Elkins et al., 2003).<br><br>32-fold increase in $K_d$ for SAM and 10-fold increase in $K_d$ for tRNA with no effect on $k_{chem}$ (Christian et al., 2016). | Unknown. | - |
| R154A | Abolishes activity (Elkins et al., 2003).<br><br>2-fold increase in $K_d$ for SAM, 15-fold increase in $K_d$ for tRNA, and 13-fold decrease in $k_{chem}$ (Christian et al., 2016).<br><br>13,000-fold reduction in $k_{cat}/K_M$ for tRNA in *Haemophilus influenzae* TrmD (Ito et al., 2015). | Unknown. | R154 forms two hydrogen bonds with guanine 37 of the tRNA substrate (Elkins et al., 2003). |
| S165L | Unknown; however, S165A substitution causes a 4-fold reduction in $k_{cat}/K_M$ for tRNA | Reduces colony size to 10% of wild-type strain in | - |

| | | | |
|---|---|---|---|
| | in *H. influenzae* TrmD (Ito et al., 2015). | *S. enterica* Typhimurium (37 °C) (Björk et al., 2001). | |
| D169A | Abolishes activity (Elkins et al., 2003).<br><br>3-fold increase in $K_d$ for SAM, 18-fold increase in $K_d$ for tRNA, and 81-fold decrease in $k_{chem}$ (Christian et al., 2016).<br><br>4100-fold reduction in $k_{cat}/K_M$ for tRNA in *H. influenzae* TrmD (Ito et al., 2015). | Unknown. | D169 is the catalytic base responsible for deprotonation of N1 in guanine 37 of the tRNA substrate (Elkins et al., 2003). |

**Supplementary Table 2. Design of *trmD* mutations at the codon level.** Mutagenic oligonucleotides were designed to introduce the indicated mutations into the *trmD* gene.

| **Substitution** | **WT codon** | **Mutant codon** | **Amino acids accessible from mutant codon by single mutation** |
|---|---|---|---|
| Y115A | TAC | GCC | Val, Asp, Gly, Thr, Pro, Ser |
| G117N | GGT | AAT | Ile, Thr, Ser, Lys, Tyr, His, Asp |
| G141A | GGT | GCT | Val, Asp, Gly, Thr, Pro, Ser |
| R154A | CGG | GCG | Val, Glu, Gly, Thr, Pro, Ser |
| S165L | TCG | CTG | Met, Val, Pro, Gln, Arg |
| D169A | GAT | GCG | Val, Glu, Gly, Thr, Pro, Ser |

**Supplementary Table 3. List of genomic mutations observed in evolved populations.** The table is derived from the output of *breseq*. The frequency of each mutation in each population is also given. The first column refers to the type of evidence used to identify the mutation: read alignment evidence (RA) or new junction evidence (JC). Mutations shown in grey were filtered from the analysis because they were considered to represent off-target mutations introduced during MAGE (mutation observed in all populations of a strain with 100% frequency), sequencing errors (mutations with a frequency <10% that were observed only near the end of reads), or mapping errors (in genes containing repetitive sequences such as *ydbA*). Mutations corresponding to the barcode sequence are not shown.

[see attached file]

**Supplementary Table 4. List of genome rearrangements observed in evolved populations.** The table is derived from the output of *breseq*. The frequency of each genome rearrangement in each population is also given. Genome rearrangements were assigned through manual annotation of new junction evidence.

[see attached file]

**Supplementary Table 5. Michaelis-Menten parameters for aminoacylation of $tRNA^{Pro}$ (CGG) by the WT and E514V variants of ProRS.** Data represent mean ± s.d. of parameters estimated from three separate experiments (except for ProRS E514V with $m^1$G37 tRNA; two separate experiments). n.d., not determined.

| tRNA | Enzyme | $k_{cat}$ ($s^{-1}$) | $K_M$ (μM) | $K_i$ (μM) | $k_{cat}/K_M$ ($M^{-1}$ $s^{-1}$) |
|---|---|---|---|---|---|
| $m^1$G37 | WT | 0.40 ± 0.07 | 0.55 ± 0.01 | 2.41 ± 0.98 | (7.31 ± 1.19) × $10^5$ |
| | E514V | 0.84 ± 0.16 | 0.27 ± 0.09 | 0.85 ± 0.15 | (3.22 ± 0.54) × $10^6$ |
| G37 | WT | n.d. | n.d. | n.d. | (2.35 ± 0.68) × $10^4$ |
| | E514V | n.d. | n.d. | n.d. | (5.22 ± 1.17) × $10^5$ |

### Supplementary Table 6. List of *E. coli* strains used in this study.

| Strain | Genotype | Source | Comments |
|---|---|---|---|
| BW25113 | | National Institute of Genetics, Japan | |
| **Barcoded strains** | | | |
| BC_S049 | BW25113; 842706_842710 delins AGCGC | MAGE | |
| BC_S050 | BW25113; 842706_842710 delins ATAGC | MAGE | |
| BC_S052 | BW25113; 842706_842710 delins AAGGC | MAGE | |
| BC_S054 | BW25113; 842706_842710 delins ACGTC | MAGE | |
| BC_S055 | BW25113; 842706_842710 delins AATAC | MAGE | |
| BC_S056 | BW25113; 842706_842710 delins AAACC | MAGE | |
| BC_S057 | BW25113; 842706_842710 delins AGTAC | MAGE | |
| ***trmD* mutants** | | | |
| BC_S089 | BC_S049; *trmD* G117N | MAGE | All *trmD* mutant strains were stored transformed with pBAD-yTrm5 and pFREE. The plasmids were cured freshly before each experiment. Y86* refers to 2,738,436_2,738,441 delins CACTAT, which gives TAA, TAG, and TGA stop codons in that order. |
| BC_S090 | BC_S050; *trmD* G141A | MAGE | |
| BC_S092 | BC_S052; *trmD* S165L | MAGE | |
| BC_S094 | BC_S054; *trmD* D169A | MAGE | |
| BC_S095 | BC_S055; *trmD* R154A | MAGE | |
| BC_S096 | BC_S056; *trmD* Y115A | MAGE | |
| BC_S145 | BC_S057; *trmD* Y86* | MAGE | |
| **Other BW25113 *trmD* wild-type derivatives** | | | |
| BC_S171 | BW25113; *proS* E19K | MAGE | |
| BC_S189 | BW25113; *proS* E19K | MAGE | |
| BC_S173 | BW25113; *proS* G200S | MAGE | |
| BC_S184 | BW25113; *proS* G200S | MAGE | |
| BC_S176 | BW25113; *proS* E514V | MAGE | |
| BC_S177 | BW25113; *proS* E514V | MAGE | |
| **BW25113 *trmD* G117N derivatives** | | | |
| BC_S148 | S089; *trmD* c(-16)t | Isolated from population G117N-2-15 | |
| BC_S150 | S089; *trmD* c(-16)t | Isolated from population G117N-2-15 | |
| BC_S147 | S089; *trmD* c(-15)t | Isolated from population G117N-1-15 | |
| BC_S152 | S089; *trmD* g(-7)c | Isolated from population G117N-2-15 | |
| BC_S155 | S089; *trmD* g(-7)c | Isolated from population G117N-2-15 | |
| BC_S149 | S089; *trmD* c(-5)a | Isolated from population G117N-2-15 | |
| BC_S151 | S089; *trmD* L61V | Isolated from population G117N-2-15 | |
| BC_S153 | S089; *trmD* L61V | Isolated from population G117N-2-15 | |

| **BW25113 *trmD* R154A derivatives** | | |
|---|---|---|
| BC_S161 | S095; *proS* E19K | Isolated from population R154A-2-22 |
| BC_S164 | S095; *proS* E19K | Isolated from population R154A-2-9 |
| BC_S167 | S095; *proS* E19K | Isolated from population R154A-2-9 |
| BC_S159 | S095; *proS* E19K, *argX* g(-5)a | Isolated from population R154A-2-22 |
| BC_S160 | S095; *proS* E19K, *argX* g(-5)a | Isolated from population R154A-2-22 |
| BC_S162 | S095; *proS* E19K, *argX* g(-5)a | Isolated from population R154A-2-22 |
| BC_S165 | S095; *proS* Y89N | Isolated from population R154A-2-9 |
| BC_S166 | S095; *proS* Y89N | Isolated from population R154A-2-9 |
| BC_S168 | S095; *proS* G107C | Isolated from population R154A-2-9 |
| BC_S169 | S095; *proS* G107C | Isolated from population R154A-2-9 |
| BC_S156 | S095; *proS* G200S | Isolated from population R154A-1-9 |
| BC_S158 | S095; *proS* G200S | Isolated from population R154A-1-9 |
| BC_S157 | S095; *proS* A231E | Isolated from population R154A-1-9 |
| BC_S163 | S095; *proS* A231E | Isolated from population R154A-1-9 |
| BC_S170 | S095; *proS* E514V | Isolated from population R154A-3-9 |
| BC_S178 | S095; *argV* t35g | MAGE |
| BC_S179 | S095; *argV* t35g | MAGE |
| BC_S181 | S095; *argX* g(-5)a | MAGE |
| BC_S209 | S095; *argX* g(-5)a | MAGE |

**Supplementary Table 7. List of oligonucleotides used in this study.**

[see attached file]
