## Supplementary figures and images for "Evolutionary repair reveals an unexpected role of the tRNA modification m^1^G37 in aminoacylation"

### Fig_5b_raw.tif

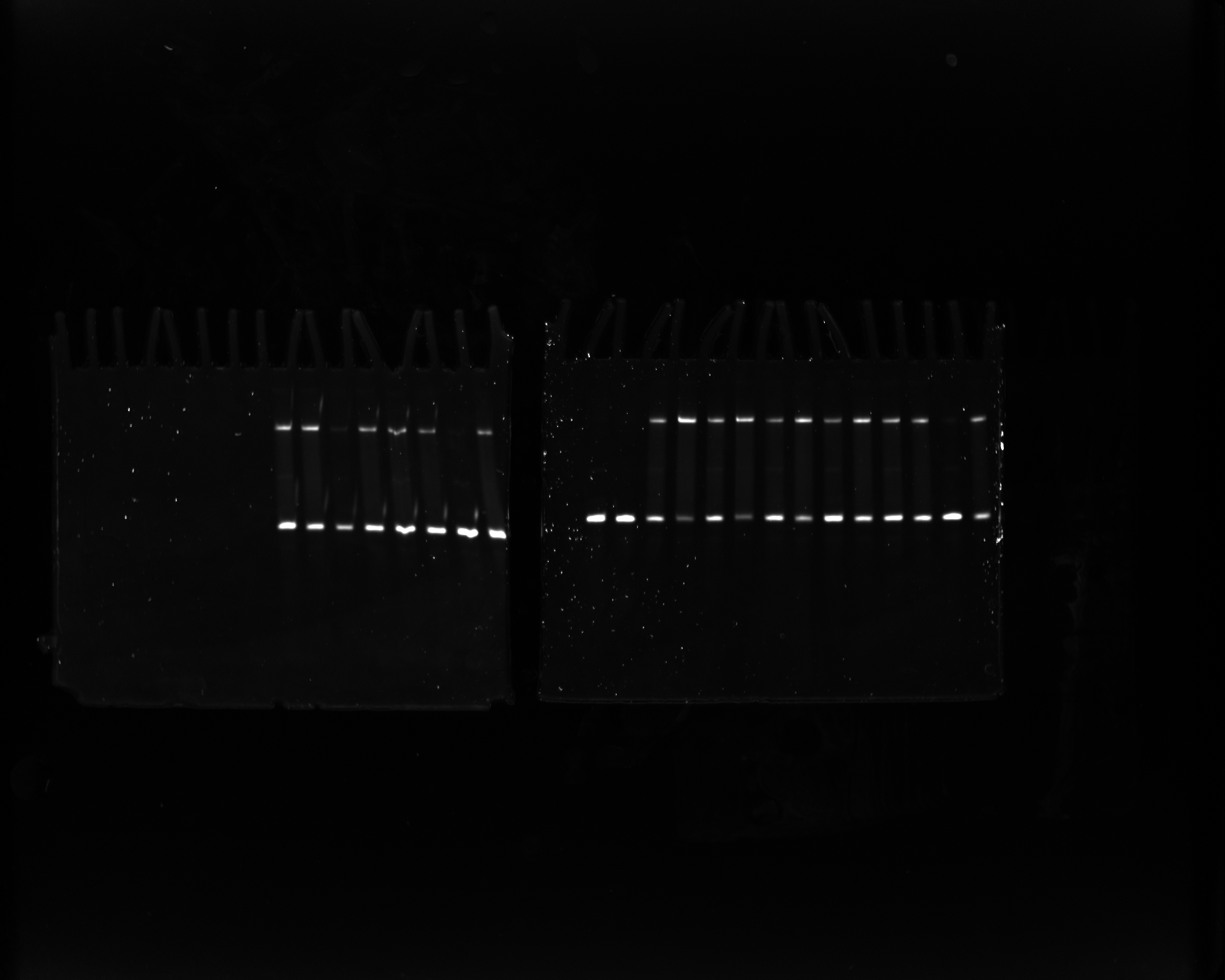

### Fig_S8a_raw.tif

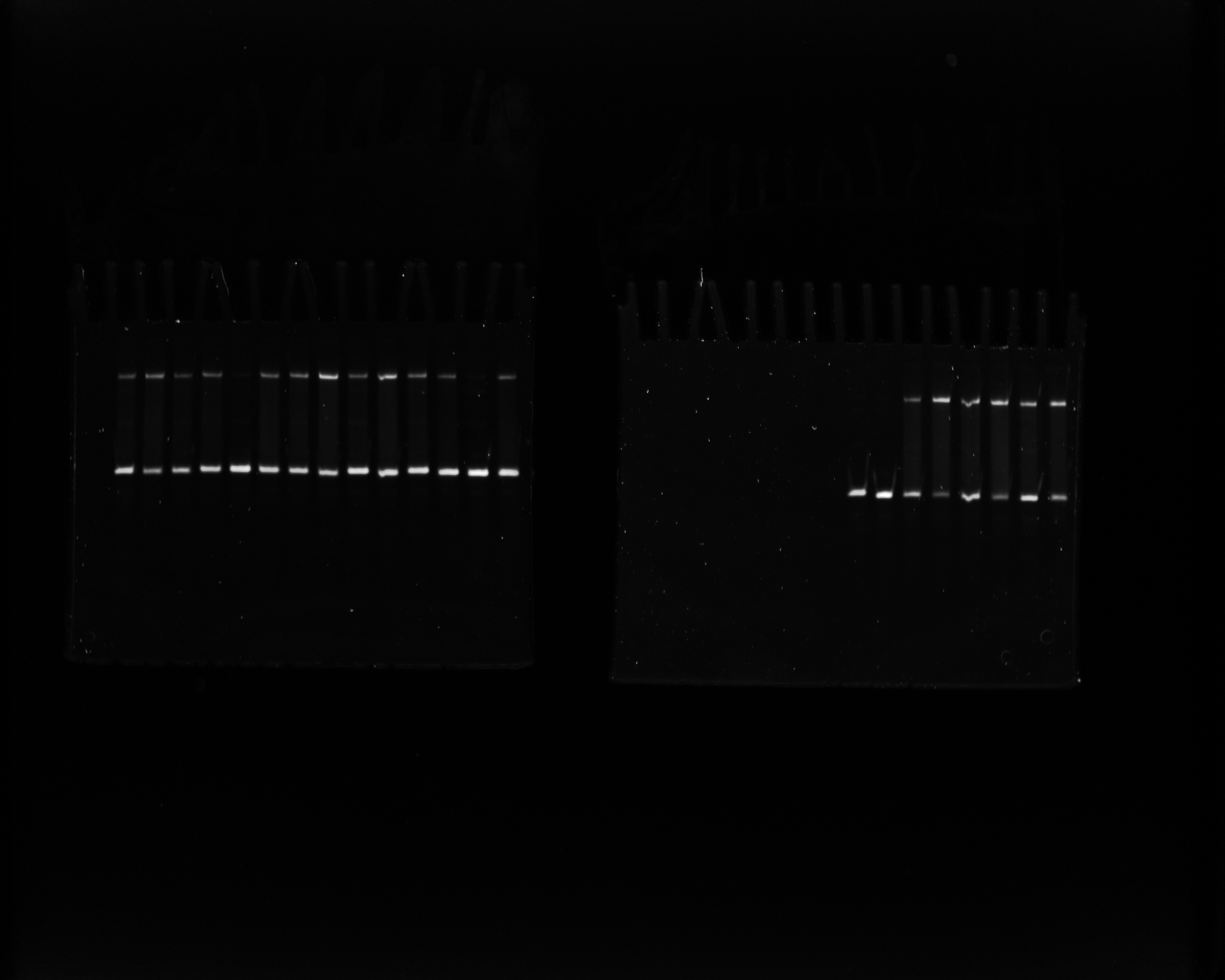

### Fig_S8b_raw.tif

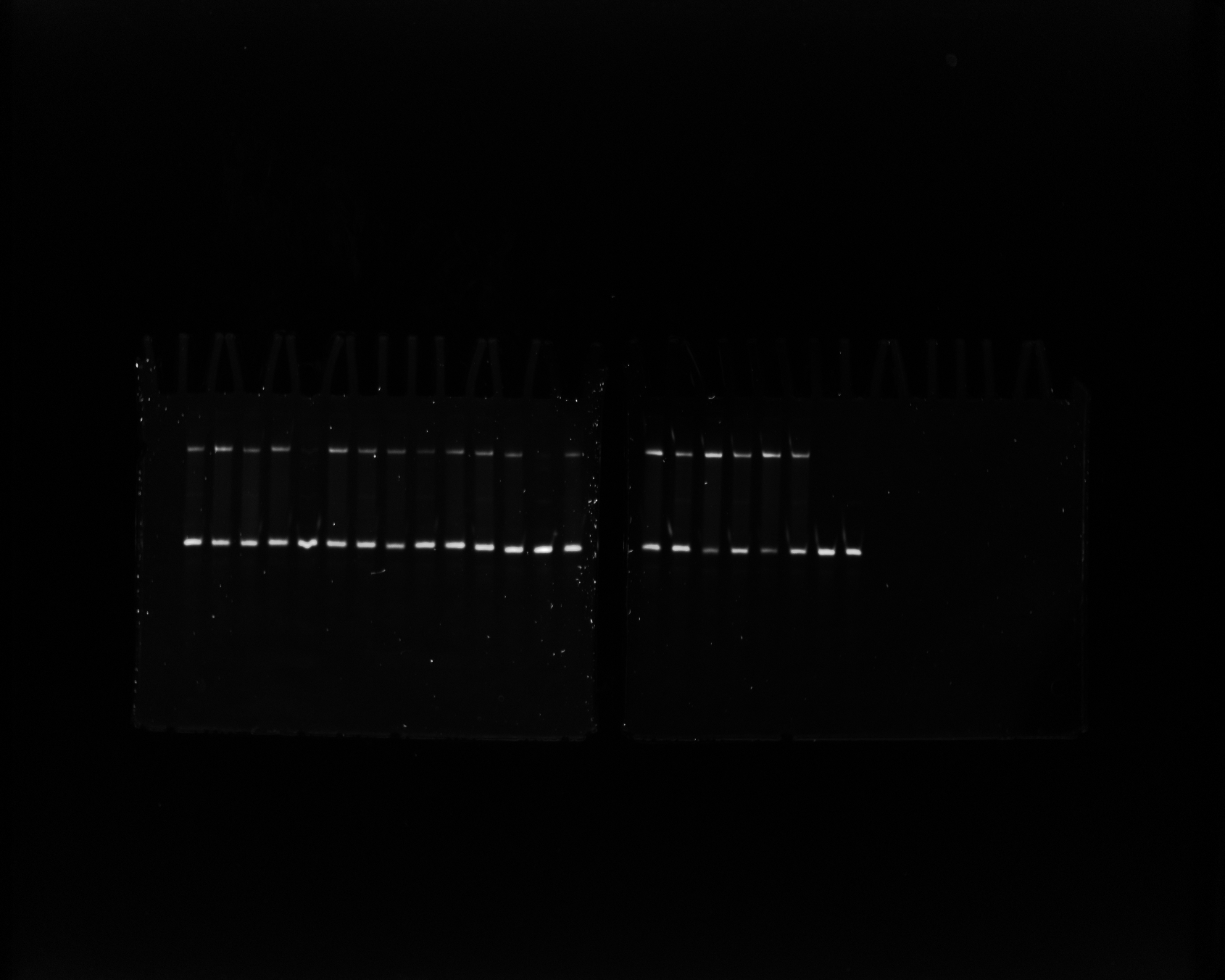

### Fig_S8c_raw.tif

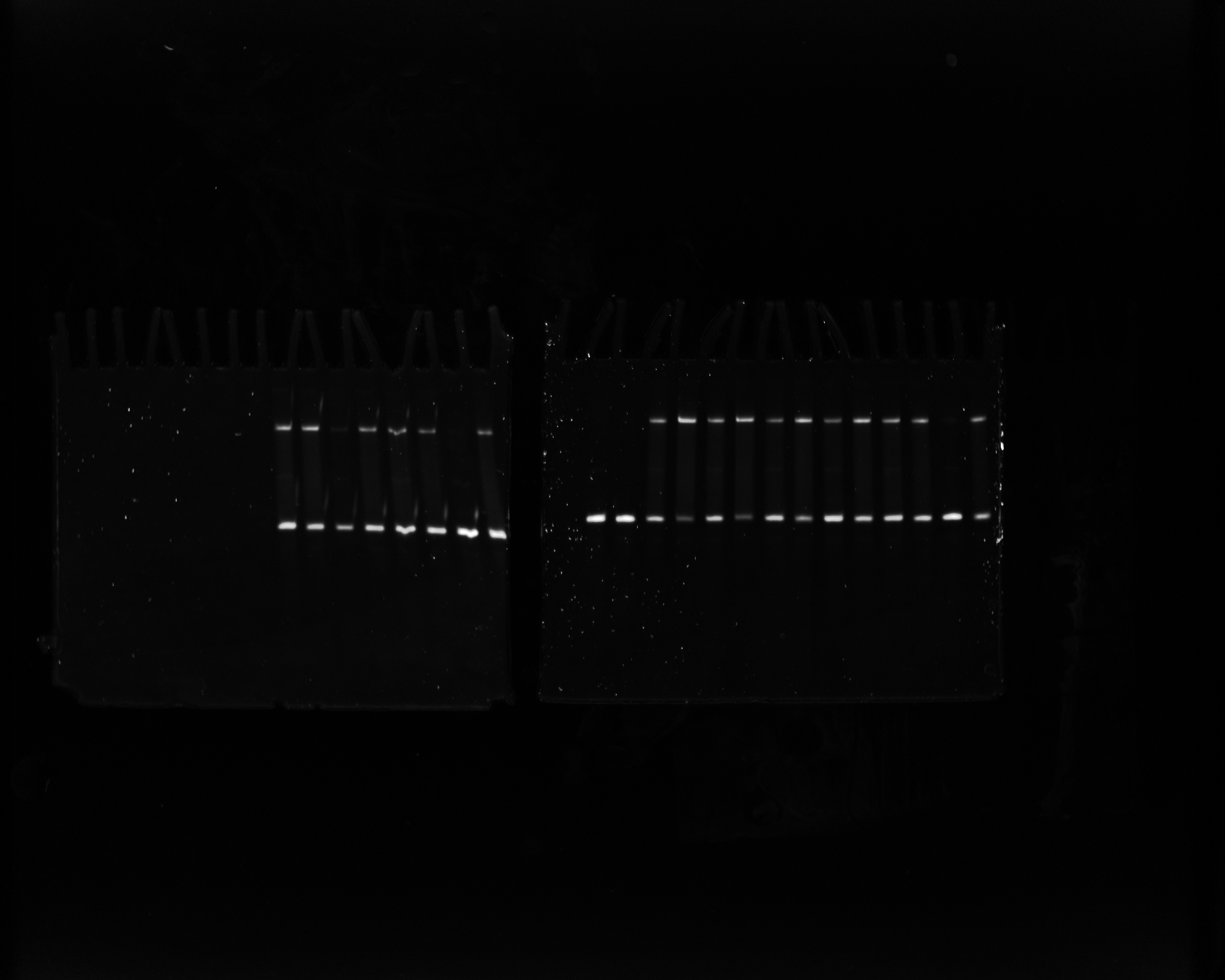

### Fig_S9a_raw.tif

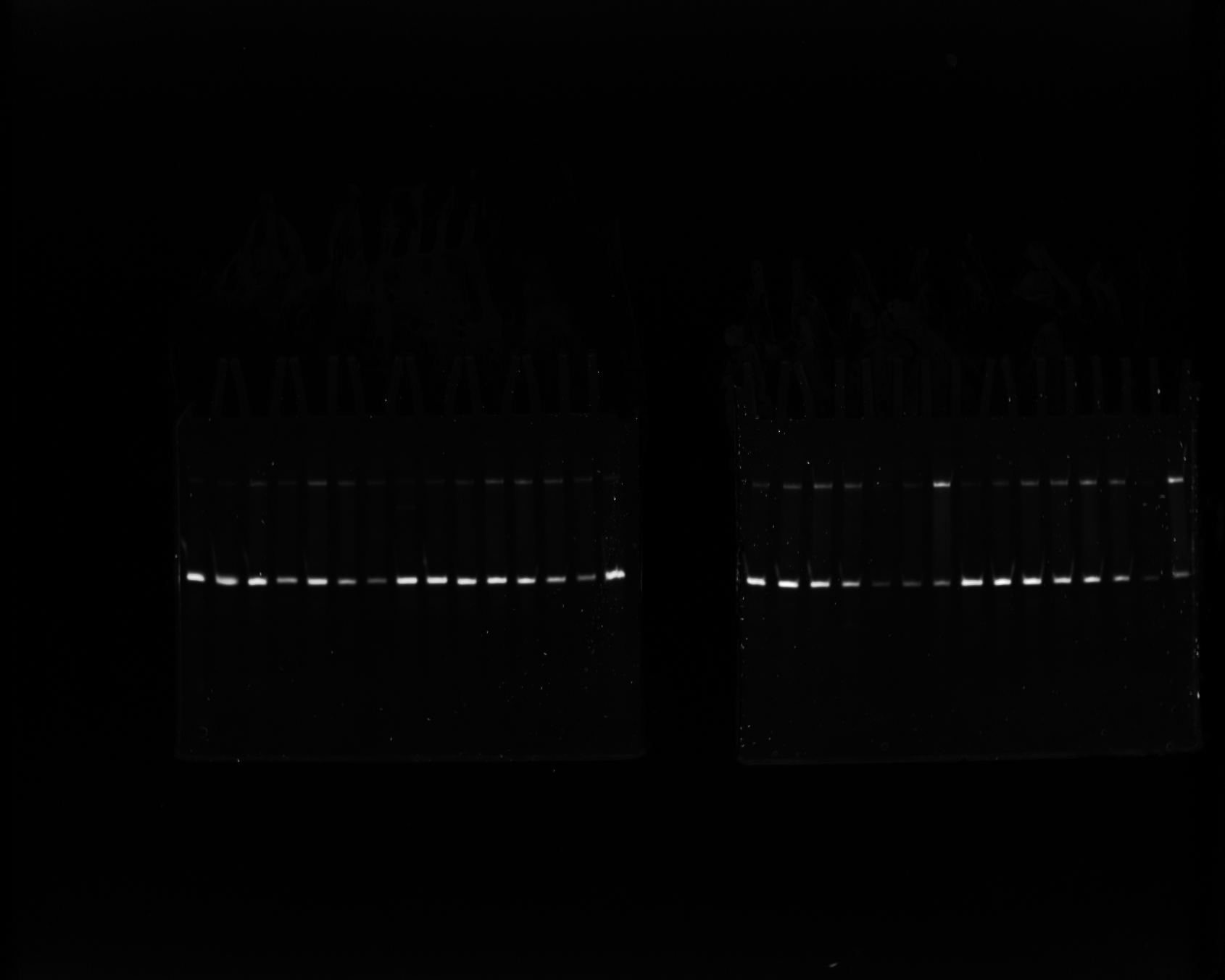

### Fig_S9b_raw.tif

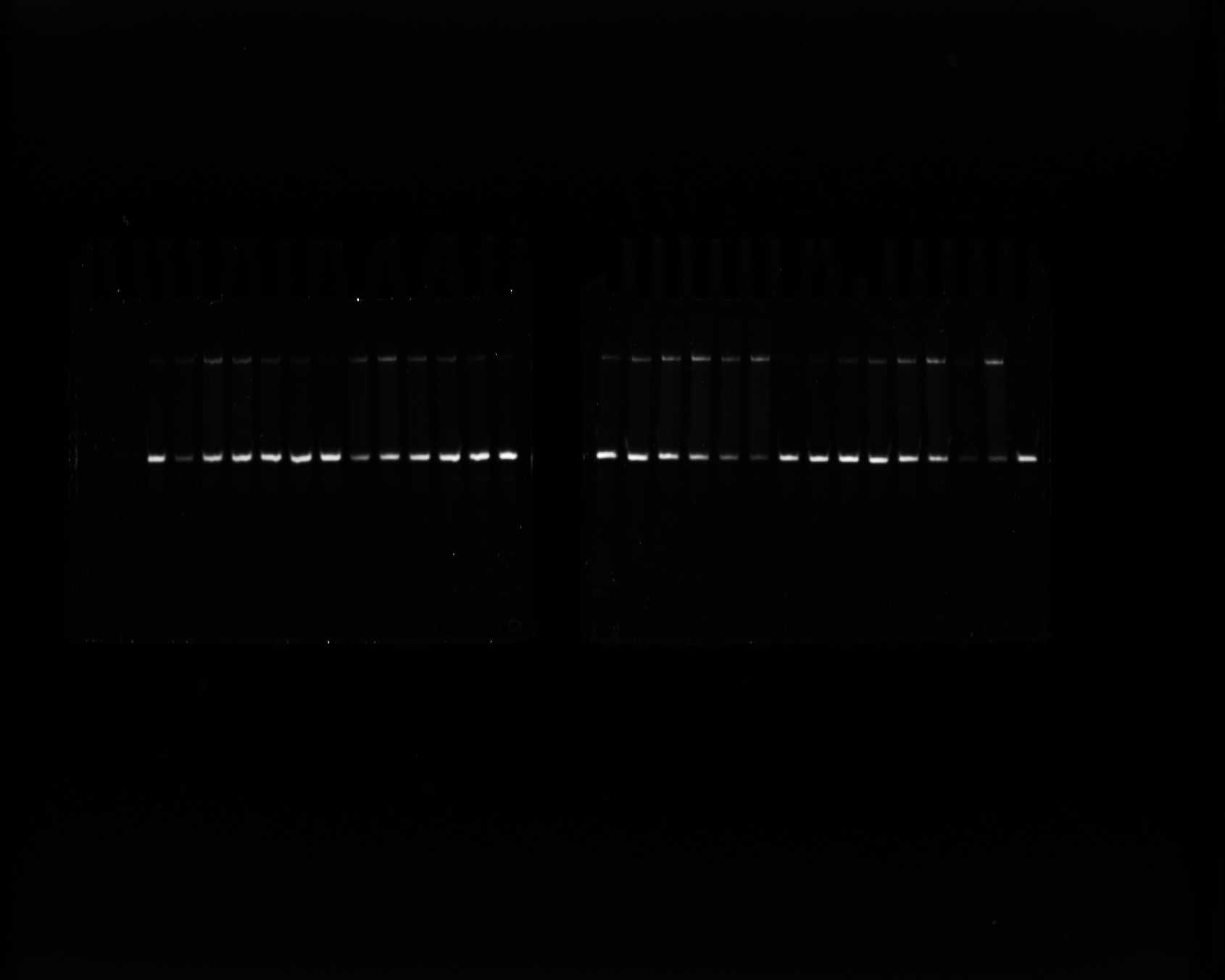

### Fig_S9c_raw.tif

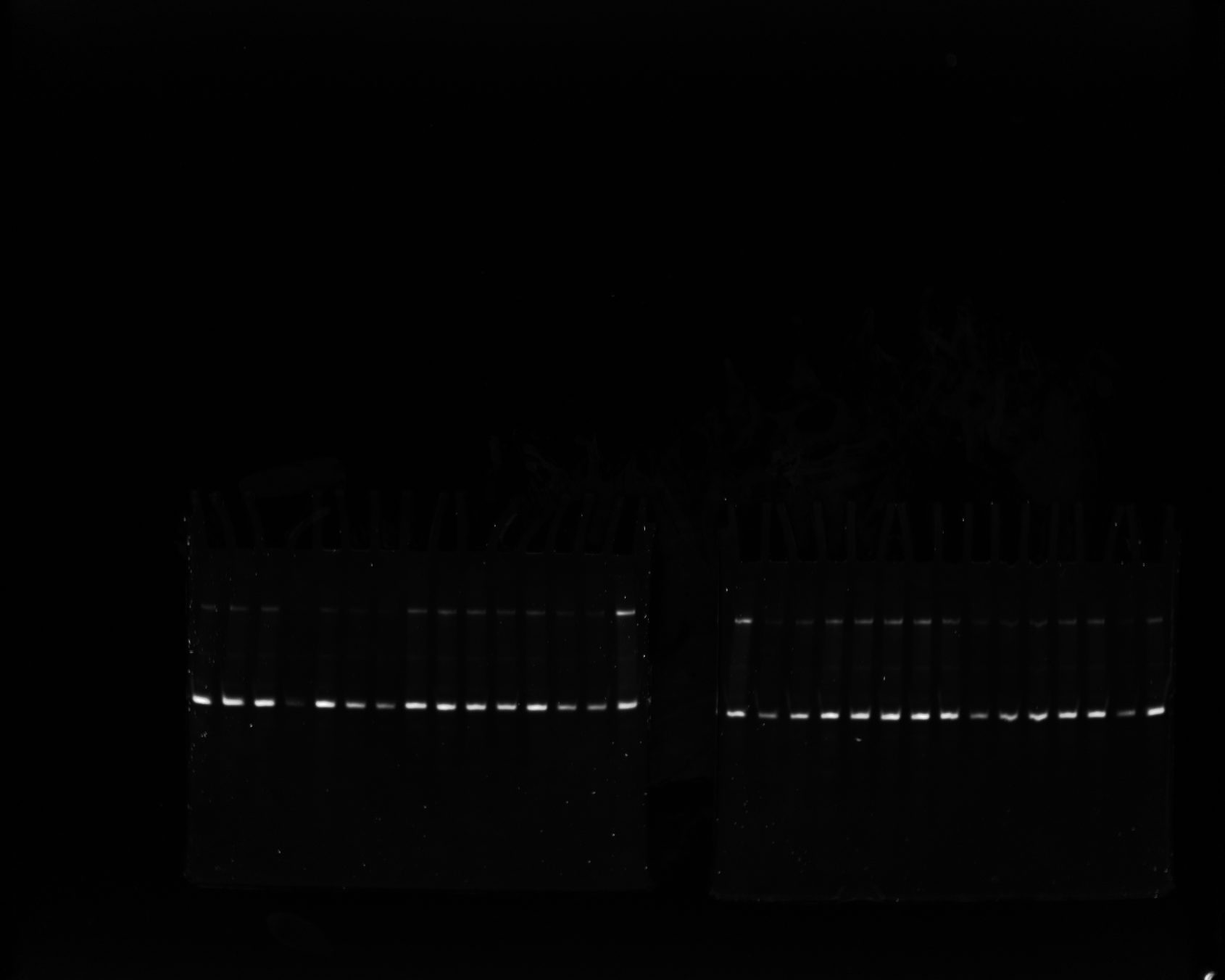
